# Hybrid genome-scale modeling and machine learning reveal cost-efficient strategies for phototrophic PHB production in *Rhodopseudomonas palustris*

**DOI:** 10.64898/2026.04.24.720730

**Authors:** Hector A. Hernandez-Gonzalez, Germán Buitrón

## Abstract

Plastic pollution from fossil-based materials is a major global environmental challenge. Microbial-derived bioplastics, such as polyhydroxybutyrate (PHB), offer a promising biodegradable alternative. However, the high substrate and operational costs of PHB production remain a major barrier to large-scale deployment. Optimizing PHB synthesis requires navigating a multidimensional design space of metabolic, nutritional, and operational variables that is impractical to explore experimentally. Here, we developed an integrated computational framework that combines a genome-scale metabolic model (GEM) of Rhodopseudomonas palustris, machine-learning surrogate modeling, Pareto multi-objective optimization, and thermodynamics-based flux analysis (TFA) to identify cost-efficient and biologically feasible PHB production strategies. Experimental and literature-derived medium compositions were translated into mechanistic constraints, enabling the GEM to generate metabolically coherent synthetic datasets that augmented sparse experimental observations. CatBoost surrogate models trained on this hybrid dataset accurately predicted PHB synthesis across thousands of hypothetical conditions, and Pareto optimization revealed operating regimes that balance PHB productivity with nutrient cost. TFA validated the thermodynamic feasibility of these strategies and refined pathway usage, reinforcing thiolase-initiated routing into PHB biosynthesis and suppressing infeasible β-oxidation–like redox loops. Overall, this hybrid GEM–ML–TFA framework identifies metabolic bottlenecks, engineering targets, and cost-optimal nutrient regimes for phototrophic PHB production, providing a scalable blueprint for rational process and strain design.

## 1. Introduction

Fossil-derived plastics remain deeply embedded in global manufacturing yet impose severe environmental and economic burdens due to their persistence and reliance on non-renewable feedstocks. Global plastic production exceeds 460 million metric tons annually, with recycling rates below 10%, resulting in widespread environmental accumulation (Geyer et al., 2017; OECD, 2022). Among sustainable alternatives, polyhydroxyalkanoates (PHAs) have emerged as promising biodegradable polymers with physicochemical properties comparable to conventional thermoplastics (Chen, 2010; Koller et al., 2017). Polyhydroxybutyrate (PHB) is a microbially synthesized polyester exhibiting high biodegradability, biocompatibility, and thermoplastic performance (Madison & Huisman, 1999). Despite decades of research, PHB remains economically constrained relative to petrochemical plastics, largely due to limitations in carbon conversion efficiency, costs, and process productivity (Choi & Lee, 1999; Dietrich et al., 2017).

Improving the economic viability of PHB production requires optimizing multiple interacting operational variables, including substrate composition, nitrogen availability, illumination intensity, and cultivation regime, that collectively govern carbon allocation, ATP generation, and intracellular redox balance. In phototrophic systems in particular, polymer accumulation is tightly coupled to the cellular requirement to dissipate excess reducing equivalents under nutrient-limited conditions. These dependencies create nonlinear trade-offs between growth and storage polymer synthesis across a multidimensional operational landscape. Systematic experimental exploration of this design space is labor-intensive, costly, and often infeasible, especially when substrate mixtures and dynamic light regimes are considered simultaneously.

Purple non-sulfur bacteria (PNSB), especially *Rhodopseudomonas palustris*, represent attractive hosts for sustainable PHB production due to their metabolic versatility and ability to grow photoheterotrophically on diverse organic substrates, including low-value waste streams (Larimer et al., 2004; McKinlay & Harwood, 2010). Their flexible redox metabolism and capacity for anaerobic phototrophic growth offer unique opportunities to integrate waste valorization with biopolymer synthesis (Capson-Tojo et al., 2020). However, this same metabolic flexibility increases system complexity: PHB accumulation arises from the interplay among substrate reduction state, nitrogen limitation, photophosphorylation efficiency, and redox-balancing pathways (Alloul et al., 2023). Consequently, identifying economically optimal operating regimes cannot rely on one-factor-at-a-time experimentation and instead requires systems-level analysis.

Genome-scale metabolic models (GEMs) provide a mechanistic framework for interrogating such trade-offs by enforcing stoichiometric mass balance and physicochemical constraints on intracellular flux distributions (Orth et al., 2010; O’Brien et al., 2015). Constraint-based approaches have been widely used to predict metabolic phenotypes and guide strain engineering strategies (Bordbar et al., 2014). Extensions incorporating thermodynamics-based flux analysis (TFA) further improve predictive realism by constraining reaction directionality according to Gibbs free energy changes and eliminating infeasible redox cycles (Henry et al., 2007; Salvy & Hatzimanikatis, 2020). Nevertheless, exhaustive evaluation of substrate combinations, nutrient ratios, and phototrophic conditions through repeated simulations remains computationally intensive. Moreover, classical flux balance frameworks do not directly resolve economic trade-offs between production performance and input cost.

Machine-learning (ML) approaches offer complementary scalability by learning nonlinear mappings between environmental inputs and production phenotypes, enabling rapid prediction across expansive parameter spaces (Carbonell et al., 2018; Kim et al., 2020). Hybrid strategies that combine mechanistic modeling with ML surrogates have begun to accelerate metabolic engineering workflows (Zampieri et al., 2019). However, data-driven models trained solely on experimental observations often suffer from limited coverage and may yield biochemically inconsistent predictions when extrapolating beyond measured conditions. Integrating thermodynamic feasibility with scalable predictive modeling, therefore, remains a critical challenge for rational engineering of complex phototrophic systems.

Here, we present a hybrid computational framework that integrates thermodynamically constrained genome-scale modeling, data-augmented machine learning, and multi-objective optimization to identify economically viable PHB production strategies in *R. palustris*. Experimentally derived and literature-reported cultivation conditions were used to constrain the GEM, generating metabolically coherent synthetic datasets that expand the feasible phenotypic space. These hybrid datasets were used to train CatBoost surrogate models that rapidly predict PHB synthesis across diverse substrate and nutrient combinations (Buitron et al., 2025). Bi-objective Pareto optimization was then applied to explicitly balance PHB production against nutrient-associated cost, identifying non-dominated operating strategies that reflect realistic economic trade-offs. Finally, TFA validated thermodynamic and redox feasibility, eliminating energetically inconsistent flux distributions.

This integrated GEM–ML–Pareto–TFA workflow reveals redox-limited operational regimes governing PHB accumulation under phototrophic conditions. It provides a scalable, cost-aware blueprint for rational metabolic and process design. Beyond PHB production, the approach establishes a generalizable methodology for coupling physicochemical constrained metabolic modeling with data-driven optimization in sustainable biomanufacturing systems.

## 2. Materials and methods

### 2.1 Experimental data collection and curation

Part of the experimental dataset used here was presented as a compound dataset (Buitrón et al., 2025). Data were generated from experiments using phototrophic mixed cultures obtained from activated sludge isolated from wastewater treatment plants. In these mixed cultures, *Rhodopseudomonas palustris* predominated, with a relative abundance greater than 72 %. The experiments explored the effects of independent variables associated with either medium composition or operational conditions (Supplementary Material) on PHB production (in mg/L and % w/w).

A subset of experiments with explicitly defined media conditions was created for downstream work, as this is required to accurately translate operational and medium parameters into uptake fluxes for metabolic modeling. This subset (shown in the Supplementary Material) primarily comprised experiments conducted under acetate, a mineral fraction of 1, no leuvinilic acid, 5 mg/L ferric citrate supplementation, 100% volume exchange, in the absence of bicarbonate, and in batch culture under continuous illumination. A general range of substrate concentrations was determined based on literature data as described next. These ranges were used to define genome-scale metabolic constraints and to enrich the dataset during the augmentation step.

### 2.2. Construction of a literature database

Since filtering the experiments with defined medium conditions reduced the database size (from 212 to 31), a literature database was constructed using PNSB data exposed to different carbon sources under variable conditions. This literature database (Supplementary Material) consisted of additional 73 experiments recovered from nine articles (Mukhopadhyay et al. 2013, Foong et al., 2022, Carlozzi et al., 2001, Mugnai et al., 2025, De Phillips et al., 1992, Brandl et al. (1989), Khatipov et al. (1998), Wu et al. (2012), Kim et al. (2011), Mckinlay et al. (2014), Ghimire et al. (2016), Luongo et al. (2017), Montiel-Corona et al. (2025)). The purpose of the literature database was to provide additional context and to expand the overall range of MCAA concentrations identified in the previous reports (Buitrón et al., 2025). Together, the experimental and literature-derived medium conditions were used to further constrain the genome-scale metabolic model (e.g., carbon substrates, NH₄⁺, and HCO₃⁻ concentrations).

Accordingly, the combined literature and experimental database was used exclusively to define model constraints and generate simulation scenarios, rather than for direct condition-specific simulations.

### 2.3 Genome-scale metabolic model

Multiple genome-scale metabolic models exist for PNSB (Alsiyabi et al., 2019; Chowdhury et al., 2022; Hädicke et al., 2011; Imam et al., 2011; Imam et al., 2013; Tec-Campos et al., 2023). Considering the high relative abundance of *R. palustris* reported in Montiel-Corona and Buitrón (2025), the *R. palustris* genome-scale metabolic model Bis A53 (*i*DT1294) was selected (Tec-Campos et al., 2023). This model includes the curated metabolic pathway for PHB synthesis, and it is compatible with the COBRApy library. Photosynthetic reactions enabling photon absorption across the visible spectrum (400–700 nm) were activated in the model to simulate an environment where all wavelengths are available, as would be the case in an open raceway pond. PHB production was decoupled from growth by defining the PHB synthesis reaction as the objective function and fixing the biomass reaction to zero flux through identical lower and upper bounds. This configuration models PHB synthesis as a steady-state phase corresponding to the second stage of a two-step process, in which biomass formation occurs first, followed by PHB accumulation during the early stationary phase as reported by Anderson & Dawes (1990) and Koller (2021). Preliminary simulations showed that when the biomass reaction was constrained to non-zero values, PHB synthesis became infeasible or yielded extremely low fluxes. That confirmed that PHB accumulation in *R. palustris* requires growth-arrested or near-stationary conditions, consistent with the second-stage PHB storage phase.

### 2.4 Calculation of uptake fluxes

To translate experimental and literature-reported medium compositions into constraints compatible with constraint-based modeling, carbon, nitrogen, and mineral availability were transformed into uptake fluxes for the genome-scale metabolic model. Each experimental data point was converted independently using its reported biomass concentration, substrate availability, and nitrogen content. In contrast, literature data were used to define physiologically realistic lower and upper bounds for substrate and nitrogen uptake reactions.

It is important to distinguish between experimentally derived substrate uptake rates and calculated substrate uptake fluxes, as these quantities serve distinct conceptual roles in constraint-based analysis. Experimentally derived uptake rates rely on direct measurements of substrate consumption (e.g., acetate disappearance; Eq. A1) but were not consistently available across all data points. Therefore, these rates were not used for comparative or multi-condition modeling. Instead, substrate uptake was calculated using COD-based conversions using Eq. A2, providing a consistent representation of total carbon availability across experimental conditions.

Medium specifications of the experimental subset were converted into flux units (mmol gDW⁻¹ h⁻¹) compatible with GEM exchange reactions. Nitrogen availability was converted from mg N L⁻¹ to flux units using Eq. A2 (Supplementary Material). Substrate uptake fluxes (e.g., acetate) were calculated by converting reported chemical oxygen demand (g COD L⁻¹) into flux units using Eq. A3 (Supplementary Material). Mineral medium components were maintained at the values of the modified Rhodospirillaceae medium (Table A1 and A2) and similarly converted from g L⁻¹ to flux units.

The resulting substrate uptake flux represents a model-level abstraction of aggregate carbon-equivalent uptake and is independent of substrate identity. Accordingly, COD-based fluxes serve as normalization constraints rather than direct physiological uptake rates. This approach is appropriate for experiments with a single defined carbon source but less ideal for experiments involving mixtures of medium-chain carboxylic acids (MCCAs), which are rarely fully characterized. Nevertheless, this strategy enables consistent and comparable carbon constraints across all conditions, while allowing experimentally resolved substrate data to anchor model scaling where available. All conversion procedures and equations are provided in the Supplementary Material.

For the literature database, substrates typically reported in g L⁻¹ were converted into uptake fluxes using the same approach. For each substrate (acetate, lactate, butyrate, isobutyrate, malate, hexanoate, and octanoate), the maximum literature-reported value was used as the lower bound of the corresponding GEM uptake reaction (negative flux, indicating uptake). In contrast, the upper bound was set to zero, thereby disallowing substrate secretion.

### 2.5 Conversion of observed PHB productions into metabolic fluxes

The PHB production observed from experiments was converted from production units (mg/L) to metabolic flux units (mmol·gDW⁻¹·h⁻¹) using Eq. A4 (in Supplementary Material).

### 2.6 Constraint-based modelling using the PHB synthesis reaction as the objective function

Each experimental datapoint was simulated independently in the genome-scale metabolic model (GEM) using its corresponding reported biomass, substrate availability, and nitrogen concentration. Literature data were not simulated directly; instead, they were used to identify carbon sources commonly associated with PHB production by purple no sulfur bacteria (PNSB) and to define physiologically realistic lower and upper bounds for substrate and nitrogen uptake reactions. These bounds were used to augment the experimental dataset by exploring multiple combinations of carbon substrates and nutrient availability within literature-reported ranges.

The GEM of *Rhodopseudomonas palustris* reported by Tec-Campos et al. (2023) was used. Simulations were performed in Python 3.12.12 using the COBRApy library in Google Colaboratory (code available on GitHub). Carbon substrates commonly used in the literature for PHB production exhibit distinct degrees of reduction (γ), reflecting differences in their intrinsic electron content. The substrates considered here include acetate (γ = 4), lactate (γ = 4), butyrate (γ = 5), isobutyrate (γ = 5), malate (γ = 3.5), hexanoate (γ = 5.33), and octanoate (γ = 5.5). For each substrate, as well as for NH₄⁺ and HCO₃²⁻, uptake-only exchange reactions were defined, with the lower bound set to the maximum literature-reported uptake rate (negative flux, indicating uptake) and the upper bound fixed at zero to prevent secretion. By selecting only the minimum (zero uptake) and maximum uptake values for each component, all possible combinations of the seven substrates and two supplements were generated (2⁹ combinations), covering both single-substrate and co-feeding scenarios.

Both standard flux balance analysis (FBA) and parsimonious FBA (pFBA) were performed using the GEM under the COBRA environment (version 0.30.0). Parsimonious flux balance analysis (pFBA), which is often considered more representative of biological systems because it assumes economical enzyme usage, maximizes the selected objective while minimizing the total flux sum, resulting in sparse flux distributions. However, because PHB accumulation in purple non-sulfur bacteria (PNSB) is frequently associated with electron sinks and overflow metabolism rather than strictly economical metabolic behavior, standard FBA was also simulated. In all simulations, the PHB synthesis reaction was defined as the objective function, with its lower bound set to zero and its upper bound constrained by a theoretical stoichiometric maximum calculated using Eq. A5 (Supplementary Material).

Top reactions were identified by ranking reactions according to their absolute flux values in pFBA and FBA. Before ranking, reactions not directly relevant to pathway-level interpretation, specifically the biomass reaction and the PHB synthesis objective reaction, were excluded from the analysis. Absolute flux values were used to prevent sign-dependent cancellation effects associated with reversible reactions. Reactions were then sorted in descending order of absolute flux magnitude, and the top 10 reactions were plotted.

In a second GEM version, PHBV biosynthesis was modelled by explicitly introducing reactions for precursor formation, monomer synthesis, and copolymerization. Propionyl-CoA was implemented as a physiological precursor via a lumped ATP-dependent activation of propionate, providing a controllable entry point for PHV biosynthesis while remaining consistent with known metabolic routes. Monomer precursor synthesis was modeled following the canonical thiolase–reductase framework: propionyl-CoA and acetyl-CoA were condensed to 3-ketovaleryl-CoA, which was subsequently reduced in an NADPH-dependent reaction to form (R)-3-hydroxyvaleryl-CoA. Copolymer formation was represented by a single lumped polymerization reaction consuming fixed proportions of (R)-3-hydroxybutyryl-CoA and (R)-3-hydroxyvaleryl-CoA, assuming a constant copolymer composition (80% PHB and 20% PHV on a molar basis). This abstraction was necessary because genome-scale models do not resolve polymer structural properties such as chain length or sequence distribution. To enable steady-state flux through the pathway and avoid dead-end metabolite accumulation, a reversible PHBV sink reaction and an associated demand reaction were included to represent polymer accumulation and removal. PHBV was therefore treated as a single flux unit with fixed composition, ensuring a physiologically consistent yet computationally tractable representation of copolymer biosynthesis.

### 2.7 Machine learning predictions

CatBoost was previously identified as the best-performing machine learning algorithm for predicting bioproduct formation by purple non-sulfur bacteria (PNSB) from operational parameters, due to its strong performance with categorical data (Buitrón et al., 2025). CatBoost is a gradient boosting algorithm based on decision tree ensembles that natively handles categorical variables through ordered boosting and internal encoding, minimizing preprocessing requirements. Accordingly, CatBoost regression was used to predict PHB synthesis flux as a function of substrate availability, nutrient conditions, and operational parameters.

Three modelling strategies were evaluated: (i) models using supplied substrate, nitrogen, and bicarbonate fluxes (i.e., pFBA or FBA input bounds); (ii) models using consumed substrate fluxes (i.e., reaction fluxes obtained either from pFBA or FBA solutions); and (iii) models using a combined feature set incorporating both supplied and consumed flux information. In all cases, continuous variables describing substrate supply or uptake were complemented with categorical operational parameters, including substrate type, illumination regime, levulinic acid presence, and operation mode. This strategy was followed either after data augmentation with pFBA or FBA.

For each dataset, CatBoost models were trained using an 80/20 train–test split, with categorical variables handled natively by the algorithm. Model performance was assessed using the root mean squared error (RMSE) and coefficient of determination (R²). In addition, five-fold cross-validation was performed on the full dataset to evaluate overall predictive robustness. Following validation, models were retrained using all available data to generate PHB flux predictions across the entire condition space.

Model interpretability was evaluated using SHAP plots values computed with a tree-based explainer. SHAP summary bar plots were used to quantify the global importance of individual features. In contrast, SHAP bee swarm plots were used to visualize how feature values influenced PHB flux predictions across conditions. These analyses enabled the identification of the most influential substrates, nutrient variables, and operational parameters driving PHB production.

### 2.8 Pareto optimality

CatBoost predicted PHB fluxes were subsequently analyzed using Pareto optimality to identify trade-offs between PHB production and substrate cost. A bi-objective optimization framework was applied to simultaneously maximize predicted PHB flux while minimizing the total cost of supplied substrate. Pareto-efficient solutions were defined as non-dominated points in the objective space, representing strategies for which improvements in PHB production cannot be achieved without increasing cost.

Substrate costs were estimated from representative laboratory-scale reagent prices obtained from Sigma-Aldrich and converted to USD per mmol. Prices are used for relative comparison only and do not reflect industrial bulk pricing. Actual costs may vary depending on supplier, purity grade, purchase volume, geographic region, and time of access.

From the resulting Pareto front, the condition achieving the highest predicted PHB flux among cost-efficient strategies was selected as the optimal PHB production strategy. Pareto fronts were visualized to compare experimental-constrained and literature-constrained GEM predictions and to identify optimal operating regions balancing productivity and economic efficiency.

### 2.9 Thermodynamics-based Flux Analysis (TFA)

Flux balance analysis (FBA) may permit thermodynamically infeasible solutions, including energy-generating cycles and reactions operating without sufficient driving force. To assess the thermodynamic feasibility of FBA optimal strategies, these solutions were further evaluated using thermodynamics-based flux analysis (TFA).

Thermodynamics-based Flux Analysis (TFA) enforces thermodynamic consistency by coupling reaction directionality with Gibbs free energy changes. For each reaction, the standard Gibbs free energy change (ΔG°ᵢ) was calculated from metabolite formation energies (Eq. 1). The actual Gibbs free energy change (ΔGᵢ) was then defined as a function of ΔG°ᵢ and intracellular metabolite concentrations (Eq. 2). Thermodynamic feasibility was enforced by constraining ΔGᵢ and reaction fluxes through linear inequalities. Standard Gibbs free energies of formation were obtained from *thermodb*, a curated experimental thermodynamic database compiled by the Laboratory of Computational Systems Biotechnology (Salvy et al. 2018) and distributed with the pyTFA framework.

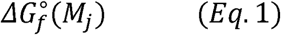

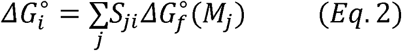

Pareto-optimal solutions were validated using thermodynamics-based flux analysis (TFA). To enable this, the *R. palustris* genome-scale metabolic model was adapted to meet pyTFA thermodynamic requirements. The modifications included updating metabolite formulas from deprotonated to protonated forms to match entries in *thermodb*. When relevant, physiological constraints from were incorporated, and all remaining reaction bounds were set to −1000 to 1000 mmol·gDW⁻¹·h⁻¹. The thermodynamically constrained model was then assembled using the standard pyTFA workflow.

Because TFA is computationally demanding, it was not applied to all simulated scenarios. Instead, it was used to validate the thermodynamic feasibility of key pathways underlying PHB synthesis under the nutrient-uptake conditions identified by the integrated GEM–CatBoost–Pareto approach. In particular, the analysis focused on reactions catalyzed by acetyl-CoA C-acetyltransferase, acetyl-CoA:acetoacetyl-CoA transferase, acetoacetyl-CoA reductase, 3-hydroxyacyl-CoA dehydrogenase (acetoacetyl-CoA), and 3-ketoacyl-CoA thiolase. The resulting model was optimized to maximize PHB synthesis. To ensure the solver did not revert to standard FBA, TFA execution was verified by confirming the activation of thermodynamic constraints and binary directionality variables. TFA flux distributions were then compared with those from standard FBA to assess the influence of thermodynamic constraints on PHB biosynthetic routing. All thermodynamic analyses were performed using pyTFA (Salvy et al., 2019).

Finally, because PHBV and other polyhydroxyalkanoates cannot be represented as single metabolites due to variable polymer length and composition, TFA was applied only to evaluate the feasibility of precursor pathways and reactions upstream of PHB synthesis rather than the polymerization reaction itself.

## 3. Results

### 3.1 Base GEM predictions

As an initial test, the GEM predicted a PHB yield (expressed as the PHB flux envelope as a function of substrate uptake) that was approximately twice the PHB flux measured for the corresponding experimental data point (Figure 1). Plotting PHB synthesis as a function of acetate uptake (Figure 1) revealed a linear increase in PHB flux with increasing acetate uptake, a pattern expected from linear programming–based methods such as flux balance analysis (FBA), which forms the basis of GEM predictions.

**Figure 1.**
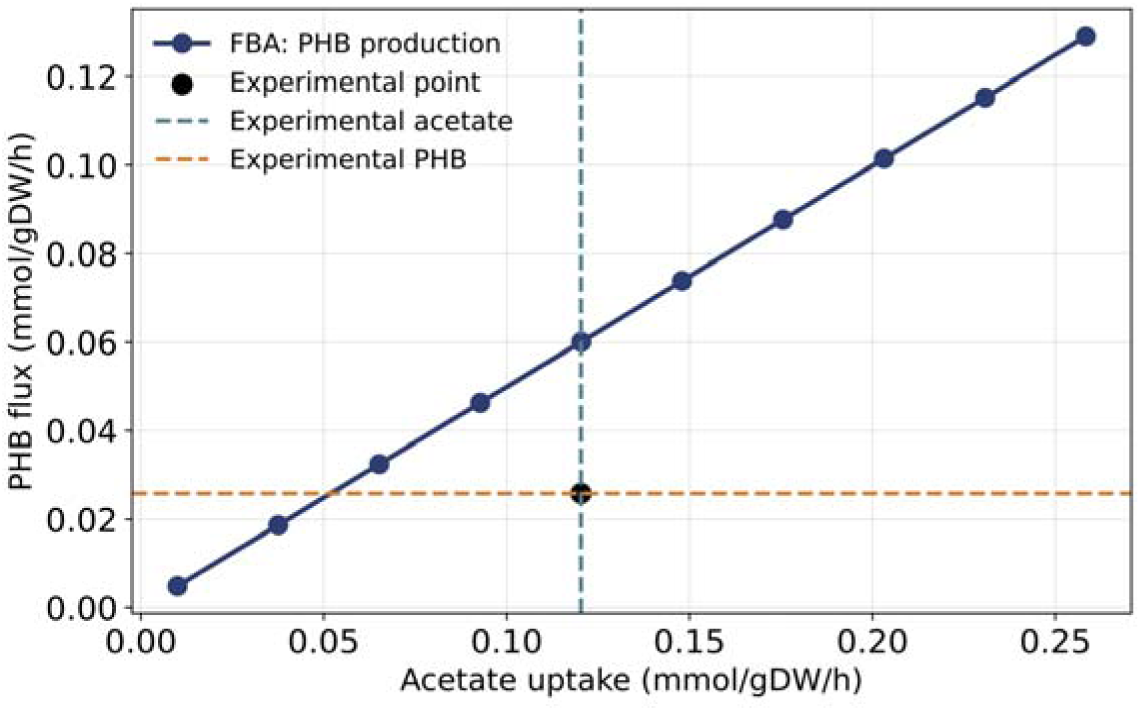
Flux balance analysis using a genome-scale metabolic model predicts a linear increase in PHB flux with acetate uptake, yielding values approximately twice those observed experimentally.

### 3.2 Data integration and augmentation

Experimental and literature data were successfully integrated into a unified dataset comprising both empirical measurements and GEM-generated synthetic data (from FBA and pFBA, respectively) across diverse combinations of carbon substrates, NH₄⁺, and HCO₃⁻ concentrations (Supplementary Material). The original dataset contained 31 experimental datapoints in which only acetate was used as the carbon substrate; after metabolic-based augmentation using a GEM constrained with literature values, the dataset expanded to 543 total conditions, of which 512 were synthetic (corresponding to 2^9^ substrate combinations). The number of synthetic datapoints produced depends strongly on the number of carbon substrates or mineral components considered (e.g., NH₄⁺, HCO₃⁻) and on the number of discrete values assigned to each. As these dimensions increase, the combinatorial expansion scales polynomially, imposing practical limits on the computational time and resources required to generate GEM simulations for all possible scenarios. Due to these computational constraints, the GEM-derived results were used to enrich and augment the dataset for training CatBoost regression models, providing both operational and metabolic features to explore the parameter space more effectively.

### 3.3 Combining metabolic modeling, CatBoost, and Pareto optimality for cost-effective optimization of PHB production

CatBoost, combined with Pareto optimization, was used to identify strategies that simultaneously maximize PHB synthesis and minimize production cost (i.e., the cost of supplying carbon substrates, NH₄⁺, and HCO₃⁻). Although these components are supplied to the system, their availability does not guarantee that they are metabolically consumed to support PHB accumulation. The GEM framework enables exploration of the metabolic pathways and uptake fluxes that cells can actually use to produce PHB. To account for this distinction, CatBoost models were evaluated under three configurations: (1) using only the supplied substrate, nitrogen, and inorganic carbon fluxes; (2) using only GEM-predicted consumed fluxes; and (3) integrating both supplied and consumed fluxes. The corresponding results for each configuration are presented in Figure 2 (panels A–C). CatBoost predictions were then followed by Pareto optimization to maximize PHB synthesis flux and minimize total costs for carbon, nitrogen, and bicarbonate.

**Figure 2.**
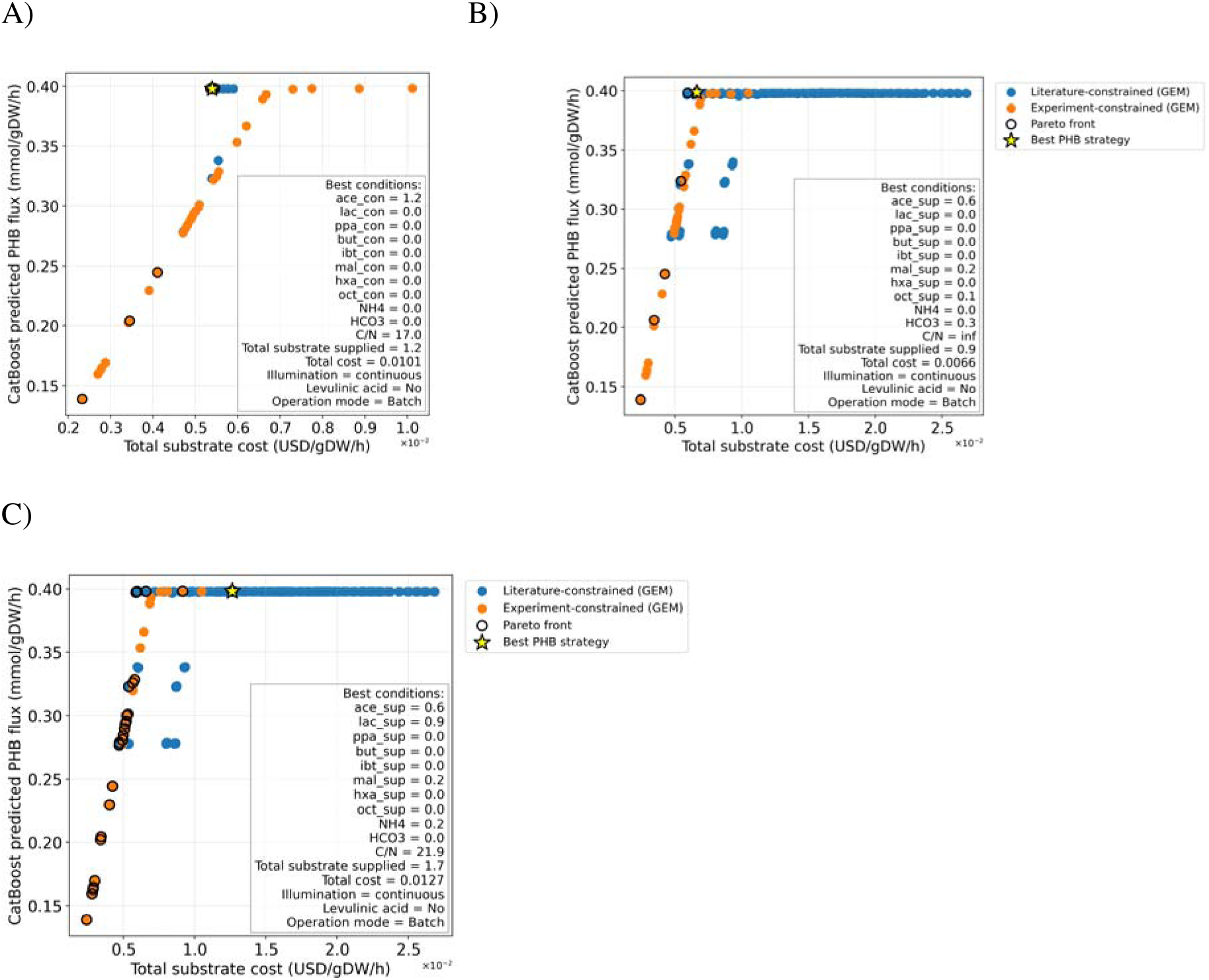
Results of CatBoost predicted and Pareto optimized PHB fluxes from FBA with different datasets (PHB synthesis upper bond = 0.39 mmol/gDW-1/hr-1). A) Only supplied nutrient fluxes are presented to CatBoost, B) Only consumed nutrient fluxes are presented to CatBoost, C) Both supplied and consumed nutrient fluxes are presented to CatBoost. The yellow star represents the Pareto optimal solution with maximal PHB production.

Several PHB experimental fluxes in Figure 2 (orange dots) reached the theoretical maximum PHB production rate, indicating that, under the model assumptions used here, these experimental conditions were already operating near their optimal metabolic potential. The solution shown in Figure 2C represents an intermediate case in which CatBoost integrates both the supplied operational conditions and the metabolically consumed fluxes, thereby capturing a balanced relationship between what is provided to the system and what the cells utilize.

Another key observation from Figure 2 is that most literature-constrained GEM predictions (blue dots) cluster at the imposed upper limit of the PHB synthesis reaction. This clustering arises because the PHB upper bound is fixed at the theoretical stoichiometric maximum calculated in Eq. (S4) (0.39 mmol gDW⁻¹ h⁻¹), a value derived from previously reported literature.

### 3.4 CatBoost Cross-validation

The Regression performance of the CatBoost models trained on the FBA augmented dataset with a PHB synthesis upper bound of 0.39 mmol gDW⁻¹ h⁻¹ (shown in Figure 2) was evaluated using five-fold cross-validation. The configuration using only supplied substrate (model “*sup”*), nitrogen, and inorganic carbon fluxes yielded an RMSE of 0.0069 ± 0.0010. The model trained using only GEM-predicted consumed fluxes during FBA achieved an RMSE of 0.0020 ± 0.0017 (model “*pfba”*). Finally, integrating both supplied and consumed fluxes during FBA resulted in an RMSE of 0.0035 ± 0.0007 (model “*sup and pfba”*). The consistently low errors and standard deviations across all three configurations indicate that the surrogate model generalizes well across the augmented dataset. Respectively, CatBoost models trained on the pFBA-augmented dataset yielded RMSE values of 0.0069 ± 0.0010, 0.0093 ± 0.0075, and 0.0076 ± 0.0053.

### 3.5 Top reactions and metabolic pathways identified from experimental data

Ranking reactions by mean absolute flux magnitude obtained from GEM simulations with experimental data revealed that the dominant flux-carrying reactions were primarily associated with light uptake and photosynthetic processes for both standard FBA and pFBA when the PHB synthesis upper bound was set to 0.3978 mmol gDW⁻¹ h⁻¹ (Figure 3). The highest-ranked reactions included photon absorption within the 680–700 nm wavelength range (*PHOA690um*), photon exchange (*EX_photon690_e*), and photosystem reactions (*PSICSum*, *PSIum*), indicating that energy acquisition and photochemical activity represent the principal drivers of the flux distribution under the simulated conditions.

**Figure 3.**
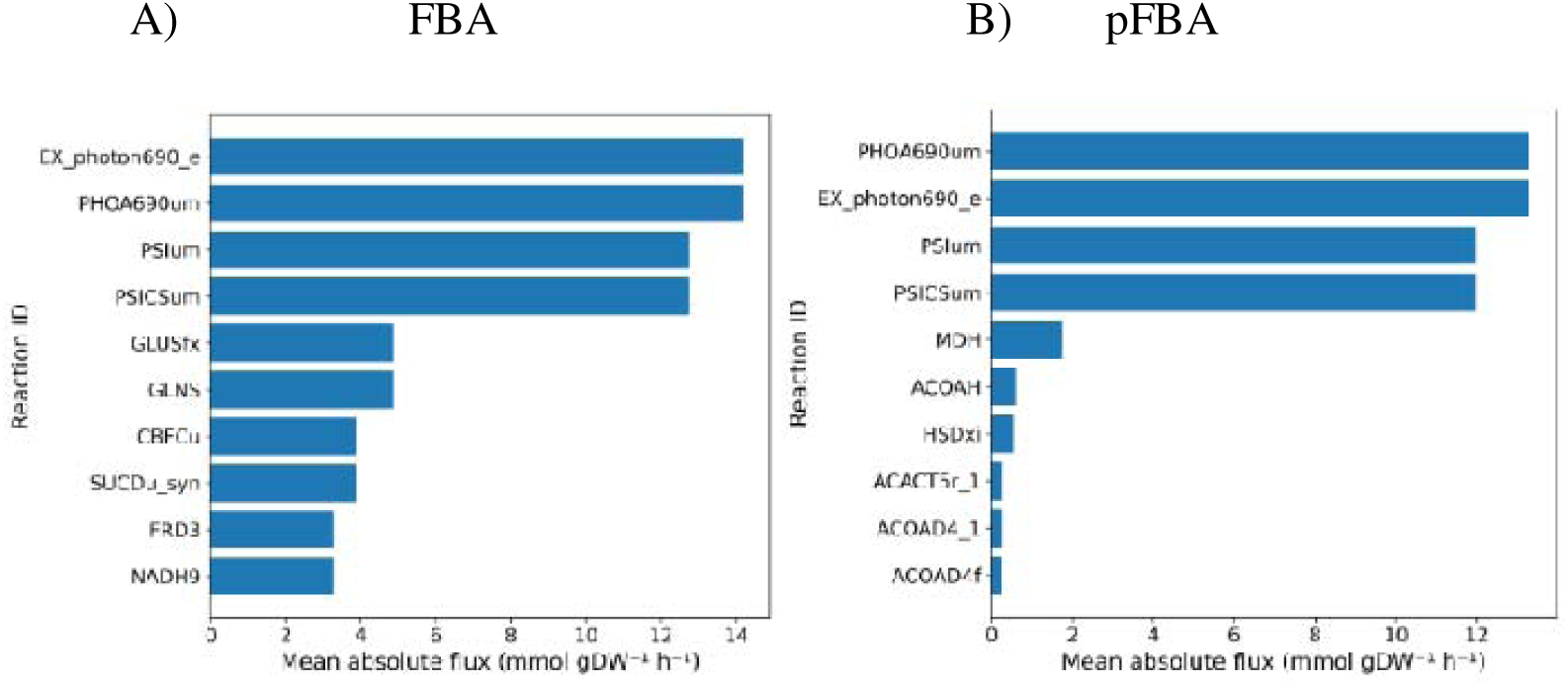
Reactions exhibiting the highest mean absolute fluxes across A) FBA and B) pFBA simulations with the PHB synthesis upper bound set to 0.3978 mmol gDW⁻¹ h⁻¹. In both cases, the dominant reactions were primarily associated with photosynthetic processes, particularly photon absorption and light harvesting within the 680–700 nm wavelength range (*PHOA690um*, *EX_photon690_e*, *PSICSum*, *PSIum*). Top flux-weighted reactions identified by pFBA predominantly belonged to central carbon metabolism (TCA cycle), amino acid biosynthesis, and fatty-acid/acyl-CoA metabolism, highlighting pathways involved in redox balance and acetyl-CoA availability relevant to PHB production. In contrast, high-flux reactions predicted by FBA were primarily associated with nitrogen, amino acid, energy, and respiration pathways.

Parsimonious FBA identified a set of flux-dominant reactions primarily associated with central carbon and acyl-CoA metabolism. Highly ranked reactions included malate dehydrogenase (MDH) and aconitate hydratase (ACOAH), indicating strong involvement of the TCA cycle. Additionally, homoserine dehydrogenase (HSDxi) contributed significantly, reflecting activity in aspartate-family amino acid biosynthesis and NADH-dependent redox processes. Reactions related to acyl-CoA metabolism, including acetyl-CoA C-acyltransferase (ACACT5r_1) and acyl-CoA dehydrogenase (ACOAD4 variants), were also prominent, suggesting active acetyl-CoA turnover and fatty-acid-associated pathways. Together, these results indicate that pFBA flux distributions are largely governed by pathways regulating energy metabolism, redox balance, and acetyl-CoA dynamics.

In addition to light-related reactions, the standard FBA identified several high-ranking reactions involved in nitrogen assimilation and amino acid metabolism. These reactions exhibited high flux relevance, including glutamine synthetase (*GLNS*), glutamate synthase (*GLUSfx*), and glutamate dehydrogenase (*GLUD_y*). Reactions associated with energy metabolism were also prominent, particularly those involved in photosynthetic electron transport (*CBFCu*) and respiratory electron transport (*SUCDu_syn*). Collectively, these results indicate that light-driven energy capture, electron transport, and nitrogen assimilation largely shape metabolic activity.

### 3.6 PHB yield in FBA is smaller and more narrowly distributed than in pFBA

PHB yield (defined as PHB flux divided by total carbon flux consumed) was compared between FBA and pFBA (Figure 4). Overall, the PHB yield predicted by FBA is lower and more narrowly distributed than that predicted by pFBA. Because both methods used the same PHB synthesis upper bound and achieved the same objective value, the higher PHB yield in pFBA can only be explained by its more efficient nutrient utilization, which uses fewer nutrients to reach the same PHB production level, thereby reducing the associated cost. The absence of an upper whisker in the pFBA boxplot shown in Figure 4 reflects a highly skewed distribution, where the maximum coincides with the upper quartile (Q3) and no values exceed the conventional 1.5×IQR whisker range. The mean overlaps with Q3, indicating that most solutions cluster near the upper bound, rendering the median marker visually indistinguishable. Overall, the median PHB yield was higher for pFBA than for FBA. A Mann–Whitney U test confirmed that the PHB yields from FBA and pFBA differ significantly, as shown in Figure 4.

**Figure 4.**
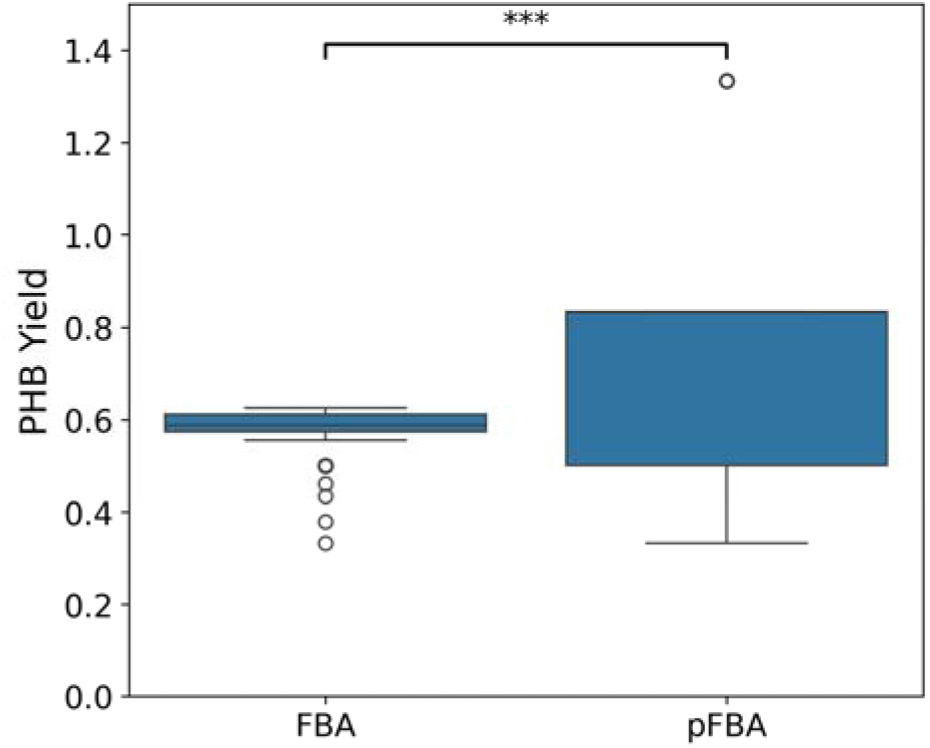
Comparison of PHB yield predicted by FBA and pFBA. PHB yield is defined as the fraction of PHB flux over total carbon flux consumed. The median PHB yield is higher in pFBA than in FBA. Statistical analysis using the Mann–Whitney U test shows a highly significant difference between the two groups (U = 97,416.5, p < 0.001, **), indicating that pFBA predictions are significantly higher than standard FBA predictions.

### 3.7 Exploring PHB synthesis beyond known physiological limits

PHB flux predictions shown in Figure 2 were inherently constrained to known metabolic limits used for the PHB synthesis reaction (0.39 mmol gDW⁻¹ h⁻¹). To explore conditions beyond reported values, the PHB synthesis upper bound was left unconstrained during FBA. This scenario was evaluated *in silico* using the same workflow, but with the PHB synthesis upper limit increased from 0.39 to 1000 mmol gDW⁻¹ h⁻¹, the maximum allowable value in the GEM. The corresponding results are shown in Figure 5 (panels A–C).

**Figure 5.**
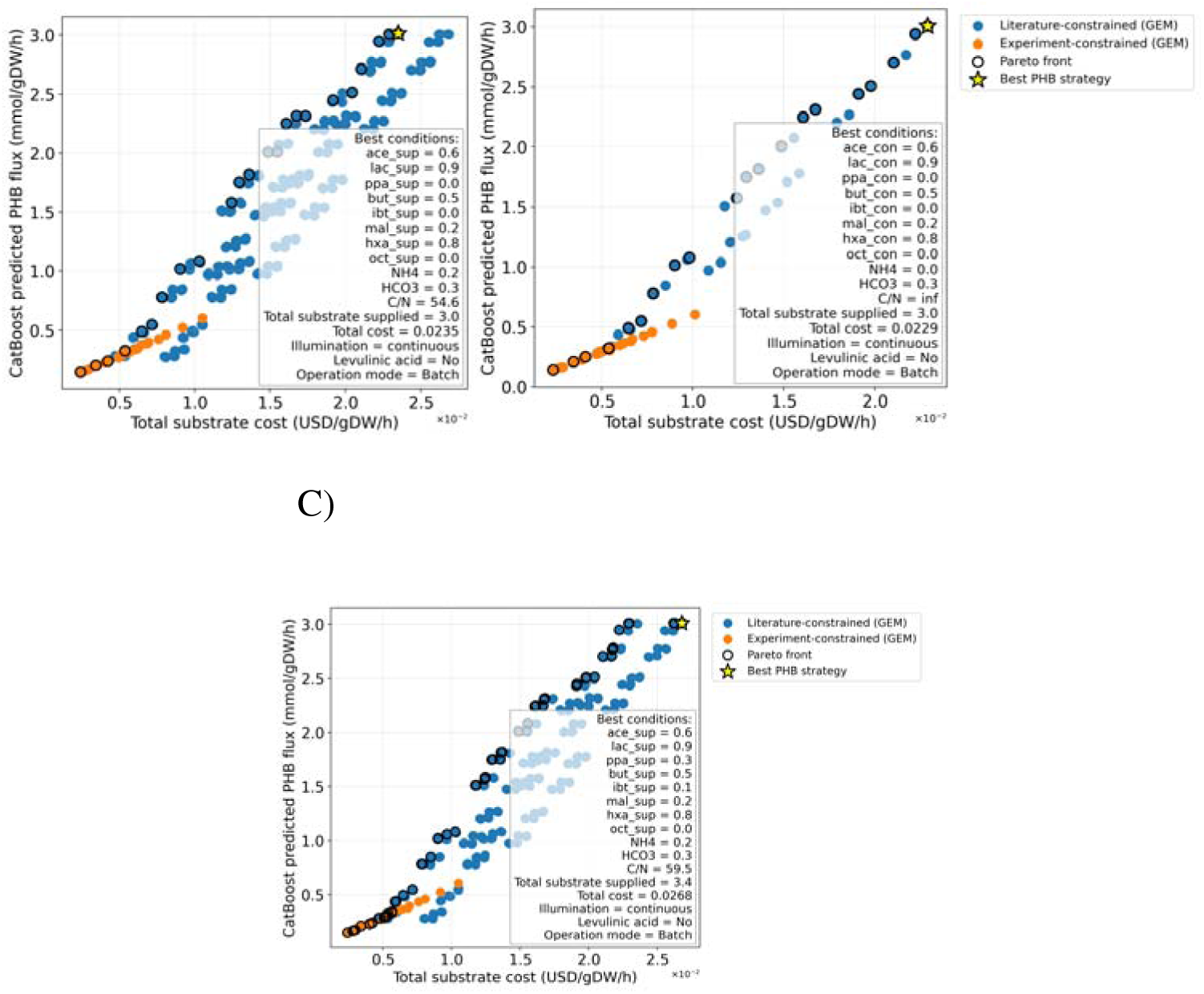
Results of CatBoost predicted and Pareto optimized PHB flux with different datasets (PHB synthesis upper bond = 1000 mmol/gDW-1/h-1). A) Only supplied substrates fluxes are presented to CatBoost, B) Only consumed substrates fluxes are presented to CatBoost, C) Both supplied and consumed substrates fluxes are presented to CatBoost. The yellow star represents the Pareto optimal solution with maximal PHB production.

Under an unconstrained scenario, the solution space expands markedly, with a few solutions reaching predicted PHB synthesis fluxes of approximately 3 mmol gDW⁻¹ h⁻¹ (Figure 5, panels A–C). In the three highest-PHB solutions, malate is among the carbon sources. Malate has been used as a pre-culture substrate for *R. palustris* (Mukhopadhyay et al., 2013) and as a carbon source for PHB production in *Rhodovulum sulfidophilum* (Foong et al., 2022) and *Cereibacter sphaeroides* (Mugnai et al., 2025). In the literature, however, malate is not considered a preferred carbon source for PHB production in *R. palustris*.

CatBoost models that incorporated consumed substrate fluxes (conditions B and C) yielded slightly higher total costs than condition A, which considered only supplied substrate fluxes. Five-fold cross-validation was also performed for this configuration. The resulting RMSE values for the three nutrient configurations were: 0.0237 ± 0.0044 when only supplied nutrient fluxes were used, 0.0032 ± 0.0016 when only consumed nutrient fluxes were considered, and 0.0035 ± 0.0007 when both supplied and consumed fluxes were included.

### 3.8 Identification of key predictors using SHAP analysis

Since 0.39 mmolgDW⁻¹h⁻¹ corresponds to the stoichiometric upper limit for PHB synthesis (Eq. S4), and the CatBoost with supplemented and consumed substrates fluxes in FBA provides a bridge between operational and metabolic capacity of PHB synthesis, this biologically grounded case was selected as the primary model for SHAP-based interpretability analysis. SHAP values reported here, therefore, reflect feature importance under physiologically plausible PHB production limits. In practice, dot colors (red/blue) in SHAP plots mark whether the feature value was high or low, respectively. The X-axis sign (positive/negative) indicates how that feature value affected the PHB prediction. Figure 6 shows that small values of total carbon consumed and their associated total costs negatively affected PHB synthesis, whereas small values of hexanoate and malate benefited PHB synthesis.

**Figure 6.**
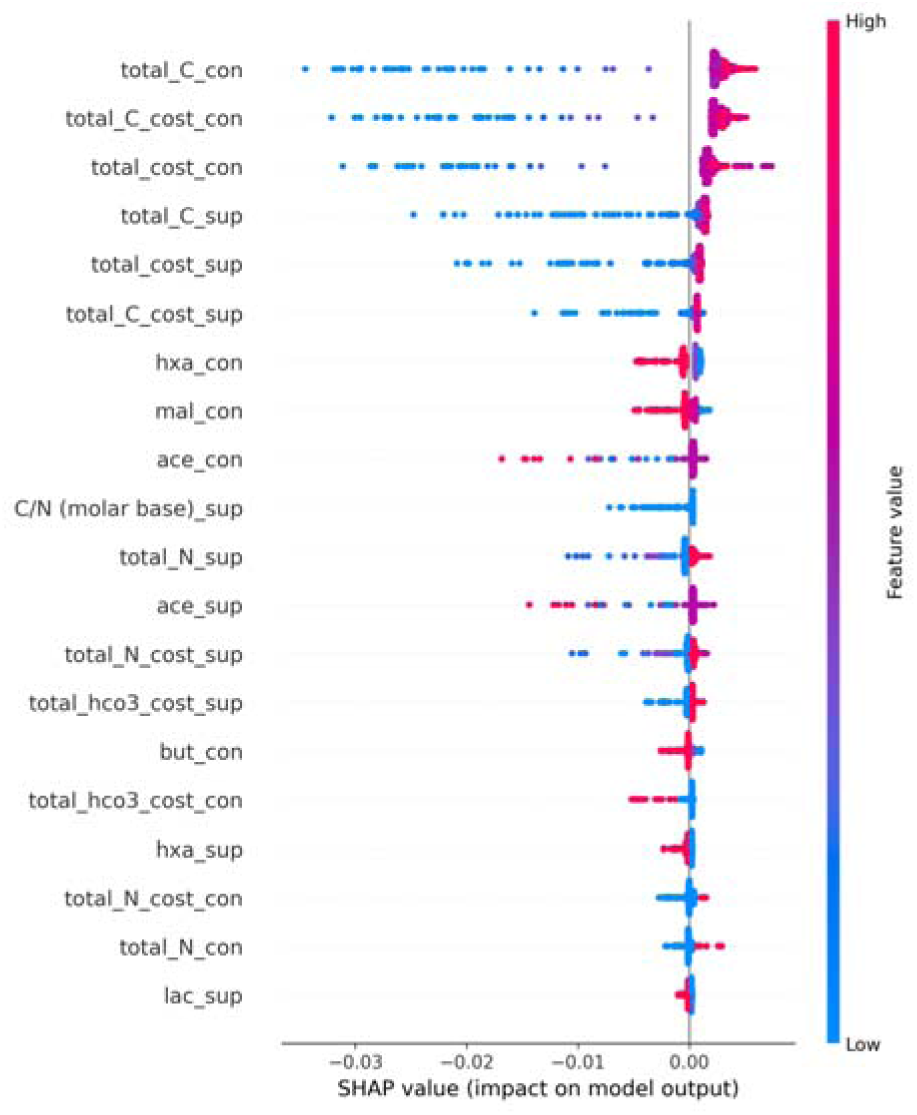
SHAP bee swarm plot of the CatBoost model constrained by the stoichiometric PHB synthesis limit (0.39 mmol gDW⁻¹ h⁻¹) in FBA. By combining supplied (sup) and consumed (con) fluxes in FBA, this model captures the link between process conditions and metabolic capacity. Dot colors denote feature magnitude, and the x-axis indicates their effect on predicted PHB flux. Features associated with high cost, high carbon supply, and low malate, hexanoate, and butyrate inputs positively influenced the model’s performance in predicting PHB flux.

An additional SHAP analysis was performed for the unconstrained model (upper bound= 1000 mmol g DW⁻¹h⁻¹) to illustrate how feature contributions shift when the PHB synthesis reaction is not restricted by known metabolic limits. Interestingly, the features with positive and negative effects on CatBoost model predictions are more strongly separated from each other in this case (see SHAP plots for the unconstrained case in the Supplementary Material).

### 3.9 Validation of Pareto-optimal solutions through thermodynamic flux analysis (TFA)

The feasibility of the five Pareto-optimal solutions derived from the CatBoost optimization strategies was evaluated using thermodynamic flux analysis (TFA). As described previously, these strategies differed in the type of training data provided to the CatBoost models. Specifically, models were trained using: (i) supplemented nutrient values (model “*sup*”), (ii) nutrient uptake fluxes obtained from pFBA (model “*pfba*”), and (iii) a combination of supplemented nutrients and pFBA uptake fluxes (model “*sup and pfba*”).

All simulations were conducted by constraining PHB synthesis (PHBS_syn) to a fixed upper bound (0.3978) while enforcing a no-growth condition (biomass flux = 0). This setup enabled a direct comparison of flux distributions predicted by FBA, pFBA, and TFA across the five optimization strategies. All approaches yielded optimal solutions, confirming the feasibility of PHB production under the evaluated conditions.

Despite their optimal solver statuses, notable differences were observed in PHB-associated reactions.

#### 3.9.1 Reaction-level differences between FBA, pFBA, and TFA reveal distinct pathway preferences

To examine how optimization assumptions shape intracellular routing toward PHB synthesis, fluxes of key reactions involved in the PHB core pathway and β-oxidation–associated dehydrogenases were compared across FBA, pFBA, and TFA using the Pareto-optimal conditions (PHB upper bound = 0.3978 mmol gDW⁻¹ h⁻¹). As shown in Table 1, the quantitative comparison, extracted from the detailed reaction-level dataset, showed that the three modeling methods differ consistently in both magnitude and prevalence of flux through central PHB precursors and competing redox-processing pathways.

**Table 1.** Quantitative comparison of flux values of seven reactions directly involved in PHB synthesis across three metabolic modeling methods (FBA, pFBA, and TFA).

| Reaction | Role | FBA | pFBA | TFA |
| --- | --- | --- | --- | --- |
| **AACOAR_syn** | Acetoacetyl-CoA reductase | 0.398 | 0.398 | 0.398 |
| **ACACT1r** | Thiolase condensation; PHB core | 0.279 | 0.279 | 0.382 |
| **HACD1** | 3-hydroxyacyl-CoA dehydrogenase<br>(acetoacetyl-CoA) | -0.119 | -0.065 | -0.016 |
| **HACD1i** | 3-hydroxyacyl-CoA dehydrogenase<br>(acetoacetyl-CoA) | 0.000 | 0.054 | 0.000 |
| **HACD1_2** | 3-hydroxyacyl-CoA dehydrogenase<br>acetoacetyl-CoA | 0.000 | 0.000 | 0.000 |
| **ACACCT** | Acetyl-CoA:acetoacetyl-CoA<br>transferase | 0.000 | 0.000 | 0.000 |
| **KAT1** | 3-ketoacyl-CoA thiolase | 0.000 | 0.000 | 0.000 |

Across FBA, pFBA, and TFA, the acetoacetyl-CoA reductase reaction (AACOAR_syn) remained fixed at the upper PHB production bound (mean ≈ 0.398 mmol gDW⁻¹ h⁻¹), confirming that this reduction step constitutes a thermodynamically and stoichiometrically robust bottleneck in PHB synthesis. This invariance indicates that all optimal solutions, regardless of the optimization formalism, must route carbon through AACOAR_syn to satisfy the imposed PHB synthesis objective.

Flux through the thiolase condensation step (ACACT1r) showed clear methodological differences. FBA and pFBA predicted similar mean flux values (≈0.279 mmol gDW⁻¹ h⁻¹), whereas TFA increased this flux substantially to ≈0.382 mmol gDW⁻¹ h⁻¹. Thermodynamic constraints, therefore, push carbon more decisively into the canonical thiolase-initiated PHB branch, consistent with previous findings showing that TFA drives ACACT1r toward the PHB upper bound under feasible ΔG conditions.

In contrast, the acetoacetyl-CoA transferase (ACACCT) and β-oxidation thiolase (KAT1) remained inactive across all methods, indicating that these alternative CoA-transfer or β-oxidation–related routes do not contribute measurably to PHB production within the Pareto-optimal space represented here.

Reactions associated with β-oxidation–like hydroxyacyl-CoA dehydrogenase activity (HACD family) showed strong method-dependent behavior. Standard FBA produced the highest HACD1 flux (mean |flux| ≈ 0.119 mmol gDW⁻¹ h⁻¹), consistent with the known tendency of FBA to accommodate broad redox-dissipating loops when no cost is associated with total flux. Parsimonious FBA reduced HACD1 flux by roughly half (mean |flux| ≈ 0.065 mmol gDW⁻¹ h⁻¹), reflecting its preference for enzyme-economical, low-flux solutions that minimize unnecessary redox cycling.

TFA further suppressed these dehydrogenase steps, driving HACD1 to near-zero mean values (≈0.016 mmol gDW⁻¹ h⁻¹) and eliminating flux entirely through HACD1i and HACD1_2. This confirms that many stoichiometrically feasible β-oxidation-related redox loops become thermodynamically infeasible once ΔG-based directionality constraints are enforced, a mechanism previously shown to prune unrealistic internal cycles in metabolic networks.

## 4. Discussion

Polyhydroxyalkanoates (PHAs) offer a compelling biodegradable alternative to petrochemical plastics, yet their high production costs remain a major barrier to deployment. Addressing this challenge requires strategies that simultaneously maximize productivity while minimizing substrate and nutrient inputs—an inherently multi-objective problem that is experimentally expensive and often intractable. In this work, we demonstrate that integrating genome-scale metabolic modeling (GEM), machine-learning surrogate modeling, Pareto multi-objective optimization, and thermodynamics-based flux analysis (TFA) creates a unified computational framework that identifies cost-efficient and biologically feasible PHB production strategies. This integration enables systematic exploration of a high-dimensional operational landscape while preserving mechanistic, metabolic, and physicochemical constraints.

### 4.1 GEM predictions and interpretation of PHB flux envelopes

Simulation of experimental conditions with the *R. palustris* GEM yielded PHB flux envelopes that scaled linearly with acetate uptake and were approximately twice the experimentally inferred PHB rates, which is an expected outcome of linear programming frameworks such as FBA (Orth et al., 2010), which capture stoichiometric upper bounds but ignore regulation, enzyme capacity, and redox constraints. These differences should not be interpreted as model failure but rather as evidence of unmodeled physiological limitations such as nitrogen availability, illumination, intracellular redox state, and stress-regulated PHB accumulation (O’Brien et al., 2015).

In *Rhodopseudomonas palustris*, PHB biosynthesis proceeds through the canonical PhaA–PhaB–PhaC pathway, where two acetyl-CoA molecules are condensed by β-ketothiolase (PhaA), reduced by acetoacetyl-CoA reductase (PhaB), and polymerized by PHB synthase (PhaC) into intracellular PHB granules (Ranaivoarisoa et al., 2019). This pathway has been identified genomically and functionally in *R. palustris* and related purple non-sulfur bacteria (Larimer et al., 2023).

### 4.2 Overflow-like behavior in FBA

In phototrophic purple non-sulfur bacteria, PHB accumulation has been described as a redox-balancing strategy under nitrogen limitation and excess-reducing conditions, serving as an intracellular electron sink that dissipates surplus NAD(P)H generated by light-driven metabolism. This redox-buffering role is particularly relevant during photoheterotrophic growth, when carbon flux, illumination intensity, and nitrogen supply collectively shape intracellular redox pressure (Alloul et al. 2023).

Standard FBA permitted substantial flux through energetically dissipative or overflow-like redox cycles, mirroring classical overflow metabolism (Basan et al., 2015), because FBA minimizes only the objective and does not penalize large, non-essential fluxes. This behavior aligns with principles of cellular resource allocation, in which metabolic flux distributions reflect trade-offs between enzyme cost and physiological benefit (Basan, 2018). In contrast, pFBA suppressed these cycles by minimizing total flux, producing enzyme-economical and physiologically coherent carbon routing and reducing power (Lewis et al., 2010). This distinction justifies the use of pFBA-derived consumed fluxes as biologically meaningful features for downstream machine-learning models.

Overall, GEM simulations provide a mechanistic context and establish an upper boundary for feasible PHB synthesis, while highlighting the need for additional layers (ML, Pareto, TFA) to reach biologically realistic and economically relevant optima.

### 4.2 Data integration and metabolic augmentation

By augmenting 31 experimental PHB-producing conditions with 512 GEM-derived synthetic scenarios, we generated a hybrid dataset that captures diverse combinations of carbon substrates, nitrogen availability, and bicarbonate inputs. This enriched design space enabled CatBoost to learn nonlinear interactions among operational parameters and PHB flux while preserving the mechanistic consistency imposed by GEM constraints. The consistently low cross-validated RMSE values indicate that CatBoost generalizes accurately across this augmented dataset.

To further assess model interpretability, reaction importance analysis identified acetoacetyl-CoA reductase (AACOAR_syn) and thiolase (ACACT1r) as dominant contributors to predicted PHB flux, along with photon uptake reactions and selected redox-linked dehydrogenases. This ranking supports the mechanistic interpretation that PHB accumulation is primarily governed by flux partitioning at the acetoacetyl-CoA branch point and by light-driven redox balance, rather than by substrate supply alone.

This hybrid augmentation approach is particularly advantageous for purple non-sulfur bacteria (PNSB), where experimental datasets are often limited and mechanistic simulations alone quickly become computationally expensive as the number of substrates and operational variables increases (Zampieri et al., 2019; Carbonell et al., 2018; Kim et al., 2020).

### 4.3 Cost-aware optimization of PHB production strategies

Coupling CatBoost predictions with Pareto multi-objective optimization allowed for simultaneous consideration of PHB productivity and substrate cost (Shoval et al., 2010). Importantly, nutrient supply does not necessarily match metabolic demand, and only models that incorporated both supplied and consumed fluxes were able to recover realistic trade-offs linking process-level inputs to cellular metabolism. These combined-feature solutions consistently lowered total costs while maintaining high PHB flux, highlighting the importance of accounting for metabolic uptake rather than relying solely on supplied nutrients.

### 4.4 Thermodynamic validation and pathway realism

Although FBA and pFBA yield stoichiometrically feasible flux states, they admit thermodynamically infeasible cycles. Applying TFA to Pareto-optimal strategies pruned these internal artifacts and reorganized flux distributions without reducing PHB productivity. TFA reinforced routing through thiolase-dependent PHB initiation and suppressed β-oxidation–like hydroxyacyl-CoA dehydrogenase loops that FBA favors but that are thermodynamically infeasible.

Photon-uptake reactions in the 680–700 nm range emerged as essential constraints on feasibility, underscoring the central role of light-driven energy metabolism in PNSB-based PHB biosynthesis.

### 4.5 Overflow suppression revisited

Notably, the overflow-like behavior seen in FBA may indicate a physiologically relevant feature of PHB accumulation in *R. palustris* (Puyol et al., 2025). FBA allows for extensive redox cycling and energy-dissipating routes that resemble the high-flux overflow metabolism PNSB use to direct excess reducing power into PHB during nitrogen limitation or redox imbalance. Overflow metabolism is often explained by proteome allocation limits that favor high-flux but less enzyme-efficient pathways under certain conditions (Basan et al., 2015). Conversely, pFBA suppresses these high-flux pathways by minimizing overall enzyme use. However, this assumption might not apply to phototrophic PHB accumulation, where redox relief rather than enzyme efficiency is the main factor. TFA further restricts only thermodynamically impossible cycles while still allowing PHB to serve as a viable overflow sink. Although both pFBA and TFA promote enzyme-efficient and thermodynamically feasible pathways, the higher-flux states predicted by standard FBA could represent redox-balancing strategies that should be tested experimentally under nitrogen-limited phototrophic conditions.

### 4.6 Engineering implications: metabolic targets and process design rules

The integrated GEM-CatBoost-Pareto-TFA workflow directly identifies actionable targets and operating principles that strengthen the metabolic engineering relevance of this study:

#### 1) AACOAR_syn as a robust metabolic bottleneck

AACOAR_syn consistently operated at the PHB upper limit (≈0.398 mmol gDW⁻¹ h⁻¹) across FBA, pFBA, and TFA, indicating an invariant bottleneck. That makes acetoacetyl-CoA reductase overexpression a high-confidence engineering strategy.

#### 2) Thermodynamic reinforcement of thiolase routing (ACACT1r)

TFA increased ACACT1r flux by ∼37%, identifying the thiolase condensation step as a thermodynamically preferred entry point into PHB synthesis. Strengthening this step via promoter engineering or cofactor optimization could enhance PHB yields.

#### 3) HACD-family dehydrogenases as targets for knockout/attenuation

HACD reactions were strongly suppressed by TFA, revealing them as futile redox sinks that dissipate reducing power away from PHB biosynthesis. Their removal could improve intracellular redox availability.

#### 4) Illumination as a critical process parameter

Photon uptake reactions (680–700 nm) dominated flux distributions and were essential under TFA, suggesting that optimizing light intensity, wavelength, and cycling regimes is a key lever for phototrophic PHB production. Notably, the experimentally inferred PHB fluxes were about half of the theoretical maximums predicted by FBA (∼0.4 mmol gDW⁻¹ h⁻¹), suggesting that targeted amplification of the bottlenecks identified above could, in theory, double polymer productivity under optimized phototrophic conditions.

### 4.7 Exploring unconstrained PHB synthesis potential

Relaxing the PHB synthesis upper bound expanded the solution space dramatically, revealing possible PHB fluxes approaching 3 mmol gDW⁻¹ h⁻¹ under hypothetical conditions. Malate emerged among the highest-performing substrates despite not being a canonical PHB precursor in *R. palustris*. This finding, supported by malate utilization in related PNSB species, warrants targeted experimental investigation. Importantly, these unconstrained solutions resulted in optimal under TFA and should therefore be validated experimentally.

### 4.8 Limitations

This study does not include gene regulatory networks, enzyme capacity constraints, dynamic resource allocation, or microbial ecology, all of which can influence PHB accumulation. Cost minimization focused only on substrate inputs rather than a comprehensive techno-economic analysis. While this framework helps identify metabolic engineering targets, it does not explicitly simulate knockout or overexpression genotypes. These limitations encourage future integration of kinetic, regulatory, and ME-model frameworks. Enzyme capacity–constrained modeling approaches (such as GECKO-type frameworks) and proteome allocation models could improve overflow predictions by explicitly considering enzyme abundance constraints, potentially aligning differences between FBA and pFBA behaviors.

### 4.9 Outlook and future directions

Future work should experimentally validate Pareto-optimal conditions, especially those involving non-canonical substrates like malate. Using regulatory FBA (Covert et al., 2001), transcriptomics-based constraints, activation of alternative electron sinks (Alloul et al., 2023), microbial ecology (Diener et al., 2020), and dynamic (Henson et al., 2014) or ME-model approaches (Oftadeh & Hatzimanikatis, 2024) could enhance predictive accuracy. Lastly, applying this framework to multi-species phototrophic consortia would allow exploration of cross-feeding, light-sharing, and resilience in outdoor systems, which are essential for scalable, low-cost PHB production.

## 5. Conclusion

In this work, we developed an integrated computational framework that combines genome-scale metabolic modeling, machine-learning surrogate modeling, Pareto multi-objective optimization, and thermodynamics-based flux analysis to identify cost-efficient and biologically feasible strategies for PHB production by *Rhodopseudomonas palustris*. By constraining the GEM with experimental and literature-derived medium compositions and augmenting these data with mechanistically consistent simulations, we generated a hybrid dataset that captures both operational variability and intracellular metabolic behavior. CatBoost surrogate models trained on this dataset accurately predicted PHB synthesis across a wide range of conditions, enabling efficient exploration of the multidimensional design space.

Pareto optimization identified operational regimes that balance PHB productivity with substrate costs, while TFA provided a mechanistic filter that eliminated thermodynamically infeasible flux patterns and refined pathway routing. Together, these components converged on a set of engineering-relevant insights, including a robust bottleneck at acetoacetyl-CoA reductase, thermodynamic reinforcement of thiolase-initiated PHB synthesis, suppression of β-oxidation-like redox sinks, and the critical role of photon uptake in the 680–700 nm range. These findings point to specific genetic and process-level targets for improving PHB production efficiency.

Overall, this hybrid GEM–ML–Pareto–TFA workflow provides a robust framework for rational bioprocess design, enabling cost-efficient optimization of phototrophic PHA production systems and supporting the development of next-generation engineered strains and cultivation methods. Besides PHB and *R. palustris*, the framework is widely applicable to other microbial hosts and bioproducts that require integrating complex metabolic, operational, and economic factors, thereby improving the predictive and translational capabilities of metabolic engineering.

