## Supplementary material for "Hybrid genome-scale modeling and machine learning reveal cost-efficient strategies for phototrophic PHB production in *Rhodopseudomonas palustris*": Supplentary material

### **Authors’ Contributions**

Conceptualization: GB and HHG, data curation: GB and HHG, formal analysis: HHG and GB, funding acquisition: GB, investigation: GB and HHG, methodology: HHG and GB, software: HHG, supervision: GB, validation: HHG, writing original draft: HHG, review and editing: GB and HHG.

#### **Calculations**

##### **Calculations for one point acetate uptake flux, (Eq. A1).**

This equation was used to calculate the acetate uptake flux for the highest PHB production reported in Montiel-Corona et al. (2022), where initial and final biomass were reported. The acetate-specific uptake rate (q_ac_) was calculated directly from experimentally measured acetate consumption, effective biomass, and cultivation time for the single reference condition. This rate represents a condition-specific physiological flux, as it is derived from the disappearance of a chemically defined substrate (e.g., acetate). The value of q_ac was used exclusively to calibrate the conversion between experimental concentration data and model flux units (mmol·gDW⁻¹·h⁻¹). Because acetate fed and residual concentrations were not available across all experimental scenarios, q_ac was not used for comparative analyses or multi-condition modeling.

$\Delta C_{ac}=C_{ac,fed}-C_{ac,remaining}$	**Eq. A1.1**

$n_{ac}=\frac{\Delta C_{ac}}{MW_{ac}}$	**Eq. A1.2**

$r_{ac}=\frac{n_{ac}}{\Delta t}$	**Eq. A1.3**

$X_{avg}=\frac{X_{f}-X_{0}}{2}$	**Eq. A1.4**

$q_{ac}=\frac{r_{ac}}{X_{avg}}=\frac{\Delta C_{ac}}{MW_{ac}\Delta tX_{avg}}(mmol{gDW}^{-1}h^{-1})$	**Eq. A1**

##### **Calculations for converting the mineral concentration in the medium to uptake flux values, Eq. (A2).**

The medium concentrations of each mineral were converted to uptake fluxes using a fixed experimental biomass of 1367 mg DW/L over 72 hrs, corresponding to the highest PHB production reported in Montiel-Corona et al., 2022. The final concentrations of minerals in the medium (µM) and the conversions to uptake fluxes are reported in Tables A1 and A2.

$n_{X}=\frac{C_{X}}{MW_{X}}$	**Eq. A2.1**

$q_{X}=\frac{n_{X}}{X_{t}\Delta t}=\frac{C_{X}}{MW_{X}X_{t}\Delta t}(mmol{gDW}^{-1}h^{-1})$	**Eq. A2.2**

where C_X_ is the final concentration of mineral X in the modified Rhodospirillaceae medium, MW_X_ is its molecular weight, X_t_ is the fixed biomass value, and t is the experiment time (in hours). Results from these formulae were used as uptake fluxes for each mineral in the GEM to simulate experimental medium conditions.

##### **Calculations for converting substrate COD into metabolic fluxes using a multipoint algorithm, Eq. (A3).**

Uptake flux calculations for multiple points in the experimental dataset were performed using a Python script that iterates over the data frame, extracts the experimental values of COD (g/L), final biomass (mg DW/L), and time (h), and applies the following formula to obtain the substrate uptake flux and the PHB production rate per row, respectively.

$q_{sub}=\frac{\frac{C_{COD}\cdot{10}^{3}}{\alpha_{COD\to sub}\cdot MW_{sub}}}{X_{t}\cdot t}$	**Eq. A3**

For acetate $\alpha_{COD\to sub}=1.067$

This formula was modified for converting the observed PHB production into PHB synthesis rate (flux).

##### **Calculations for converting PHB production into metabolic fluxes within a multipoint algorithm, Eq. (A4).**

$n_{PHB}=\frac{C_{PHB}}{MW_{PHB}}$	**Eq. A4.1**

$q_{PHB}=\frac{n_{PHB}}{X_{t}\Delta t}=\frac{C_{PHB}}{MW_{PHB}X_{t}\Delta t}(mmol{gDW}^{-1}h^{-1})$	**Eq. A4.2**

##### **Calculations of theoretical PHB synthesis stoichiometric maximum (Eq. A5).**

The maximum amount of PHB that can be produced from an organic substrate, for example acetate, is 0.39 mol PHB/mol acetate according to the following equations. This is the stoichiometric upper bound on PHB synthesis.

$q_{PHB}^{max}=Y_{{PHB}/{Ac}}^{max}\cdot q_{Ac}=\frac{2}{4}\cdot\frac{\gamma_{Ac,C}}{\gamma_{PHB,C}}\cdot q_{Ac}=\frac{2}{4}\cdot\frac{3.5}{4.5}\cdot q_{Ac}\approx0.39\cdot q_{Ac}$	**Eq. A5.1**

Here, $q_{PHB}^{max}$ denotes the maximum stoichiometric PHB synthesis rate (mmol PHB/gDW/h), $Y_{{PHB}/{Ac}}^{max}$ is the maximum stoichiometric yield of PHB on acetate (mol PHB/mol acetate). The variable $q_{Ac}$ corresponds to the specific acetate uptake rate, defined as the rate of acetate consumption normalized by the existing biomass (mmol acetate gDW/h). The variables $\gamma_{Ac,C}$ and $\gamma_{PHB,C}$ denote the degree of reduction per carbon atom in acetate and PHB, respectively, and account for redox constraints on carbon conversion. The factor $\frac{2}{4}$ normalizes carbon fluxes by accounting for the two carbon atoms per acetate molecule and the four carbon atoms per PHB monomer. Together, these terms define a stoichiometric upper bound for PHB synthesis from acetate, assuming unlimited light supply and in the absence of energetic or kinetic constraints.

Acetate uptake rates ($q_{Ac}$) reported in the literature are 2.0 ± 0.0 μmol mg$\cdot$DCW^-1^$\cdot$h^-1^ under growing conditions (McKinlay & Harwood, 2010) and 0.042 ± 0.007 μmol mg^-1^$\cdot$DCW^-1^$\cdot$h^-1^ under non-growing conditions (McKinlay et al., 2014). Because cells transitioning into nitrogen-starved, non-growing conditions are likely to have already taken up acetate at higher rates during the growth phase, an effective acetate uptake rate, corresponding to the average of these two values (i.e., 1.02 μmol mg$\cdot$DCW^-1^$\cdot$h^-1^), was used. This amount was used as the upper limit of the PHB synthesis reaction in the GEM.

$q_{PHB}^{max}\approx0.39*1.02\approx0.3978$	**Eq. A5.2**

Alternatively, the following calculation can be used when the PHB production process is modeled as a one-step growth-coupled process (which is not the case in this research). This method (shown in Eq. A5.3) is reported only as a reference and comparison with the theoretical maximum of PHB synthesis presented above.

$q_{PHB,max}=\mu_{max}\cdot f_{PHB}\cdot\frac{1000}{MW_{PHB}}$	**Eq. A5.3**

$=0.08h^{-1}\cdot0.50\cdot\frac{1000mgg^{-1}}{86.09mg{mmol}^{-1}}$	**Eq. A5.4**

$=0.4646mmol{gDW}^{-1}h^{-1}$	**Eq. A5.5**

Where, $\mu_{max}$ is the standard reported growth rate of *R. palustris* under photoheterotrophic growth $f_{PHB}$ (g PHB gDW^-1^) represents the mass fraction of PHB within newly formed biomass. The factor 1000 converts grams of PHB to milligrams, while $MW_{PHB}$ the molecular weight of the PHB monomer enables conversion from a mass-based rate to a molar flux.

#### **Table A1.** Modified Rhodospirillaceae medium used for experiments reported by Buitron et al. (2025).

| **Compound (as listed)** | **g·L⁻¹** | **Molar mass (g·mol⁻¹)** | **M (mol·L⁻¹)** | **Final (µM)** | **Reaction in GEM** | **Uptake flux (mmol·gDW⁻¹·h⁻¹)** |
| --- | --- | --- | --- | --- | --- | --- |
| **Yeast extract** | 0.30 | Variable | Variable | Variable | NA | NA |
| **Sodium acetate (NaCH₃COO)** | Experiment dependent | Experiment dependent | Experiment dependent | Experiment dependent | EX_ac_e | Experiment dependent |
| **(NH₄)-acetate (NH₄CH₃COO)** | Experiment dependent | Experiment dependent | Experiment dependent | Experiment dependent | EX_ac_e | Experiment dependent |
| **NH₄Cl** | Experiment dependent | Experiment dependent | Experiment dependent | Experiment dependent | EX_nh4_e | Experiment dependent |
| **KH₂PO₄** | 0.50 | 136.088 | 3.67×10⁻³ | 3,674 µM | EX_pi_e | 0.03733 |
| **MgSO₄ (anhydrous)** | 0.19 | 120.365 | 1.58×10⁻³ | 1,579 µM | EX_mg2_e | 0.01604 |
| **NaCl** | 0.40 | 58.44 | 6.84×10⁻³ | 6,845 µM | EX_na1_e | 0.06955 |
| **CaCl₂·2H₂O (assumed)** | 0.05 | 147.01 | 3.40×10⁻⁴ | 340.1 µM | EX_ca2_e | 0.003455 |
| **Fe(III) citrate (assumed FeC₆H₅O₇)** | 0.01 | 244.951 | 4.08×10⁻⁵ | 40.82 µM | EX_fe3_e | 0.0004147 |
| **L-cysteine·HCl (assumed)** | 0.30 | 157.614 | 1.90×10⁻³ | 1,903 µM | Reaction not available in GEM | Reaction not available in GEM |
| **Thiamine·HCl (assumed)** | 0.001 | 301.814 | 3.31×10⁻⁶ | 3.31 µM | Reaction not available in GEM | Reaction not available in GEM |
| **Nicotinic acid (C₆H₅NO₂)** | 0.001 | 123.113 | 8.12×10⁻⁶ | 8.12 µM | Reaction not available in GEM | Reaction not available in GEM |
| **Trace element solution** | 0.4 mL |  |  |  |  |  |

#### **Table A2.** Trace element solution of the modified Rhodospirillaceae medium.

| **Compound (stock)** | **g·L⁻¹ (stock)** | **Molar mass (g·mol⁻¹)** | **Stock M (mol·L⁻¹)** | **Final in medium (mol·L⁻¹)** | **Final in medium (µM)** | **Reaction in GEM** | **Uptake flux (mmol·gDW⁻¹·h⁻¹)** |
| --- | --- | --- | --- | --- | --- | --- | --- |
| **ZnSO₄·0.7H₂O** | 0.10 | 174.051 | 5.745×10⁻⁴ | 2.298×10⁻⁷ | 0.230 µM | EX_zn2_e | 0.000002337 |
| **MnCl₂·4H₂O** | 0.03 | 197.904 | 1.516×10⁻⁴ | 6.064×10⁻⁸ | 0.061 µM | EX_mn2_e | 0.000000620 |
| **H₃BO₃** | 0.30 | 61.834 | 4.852×10⁻³ | 1.941×10⁻⁶ | 1.941 µM | EX_bo3_e | 0.00001972 |
| **CoCl₂·6H₂O** | 0.20 | 237.926 | 8.406×10⁻⁴ | 3.362×10⁻⁷ | 0.336 µM | Reaction not available in GEM | 0.000003414 |
| **CuCl₂·2H₂O** | 0.01 | 170.482 | 5.866×10⁻⁵ | 2.346×10⁻⁸ | 0.023 µM | EX_cu2_e | 0.000000234 |
| **NiCl₂·6H₂O** | 0.02 | 237.686 | 8.414×10⁻⁵ | 3.366×10⁻⁸ | 0.034 µM | EX_ni2_e | 0.000000345 |
| **NaMoO₄·2H₂O** | 0.03 | 218.972 | 1.370×10⁻⁴ | 5.480×10⁻⁸ | 0.055 µM | EX_mobd_e | 0.000000559 |

In addition to the previous specification of Rhodospirillaceae medium, the following specifications were used for CO2, O2, and HCO3, as summarized in Table A3.

**Table A3.** Additional constraints used in the genome-scale metabolic model.

| Compound (as listed) | Reaction name in GEM | Uptake flux (mmol·gDW⁻¹·h⁻¹) |
| --- | --- | --- |
| CO2 | EX_co2_e | 0.0 |
| O2 | EX_co2_e | 0.0 |
| HCO3 | EX_hco3_e | Dependent on experiment |

### **7. Supplementary Results**

#### CatBoost and Pareto optimization in pFBA

As a complementary approach to FBA, Parsimonious FBA (pFBA) finds the solution that maximizes the objective (PHB synthesis) while minimizing the total sum of flux. In other words, it yields a parsimonious flux that minimizes enzyme usage. CatBoost and Pareto optimization were performed with an upper bound of 0.398 mmol gDW⁻¹ h⁻¹ for PHB synthesis in pFBA. The three scenarios were optimal in TFA validation, indicating thermodynamic feasibility.

A)	B)

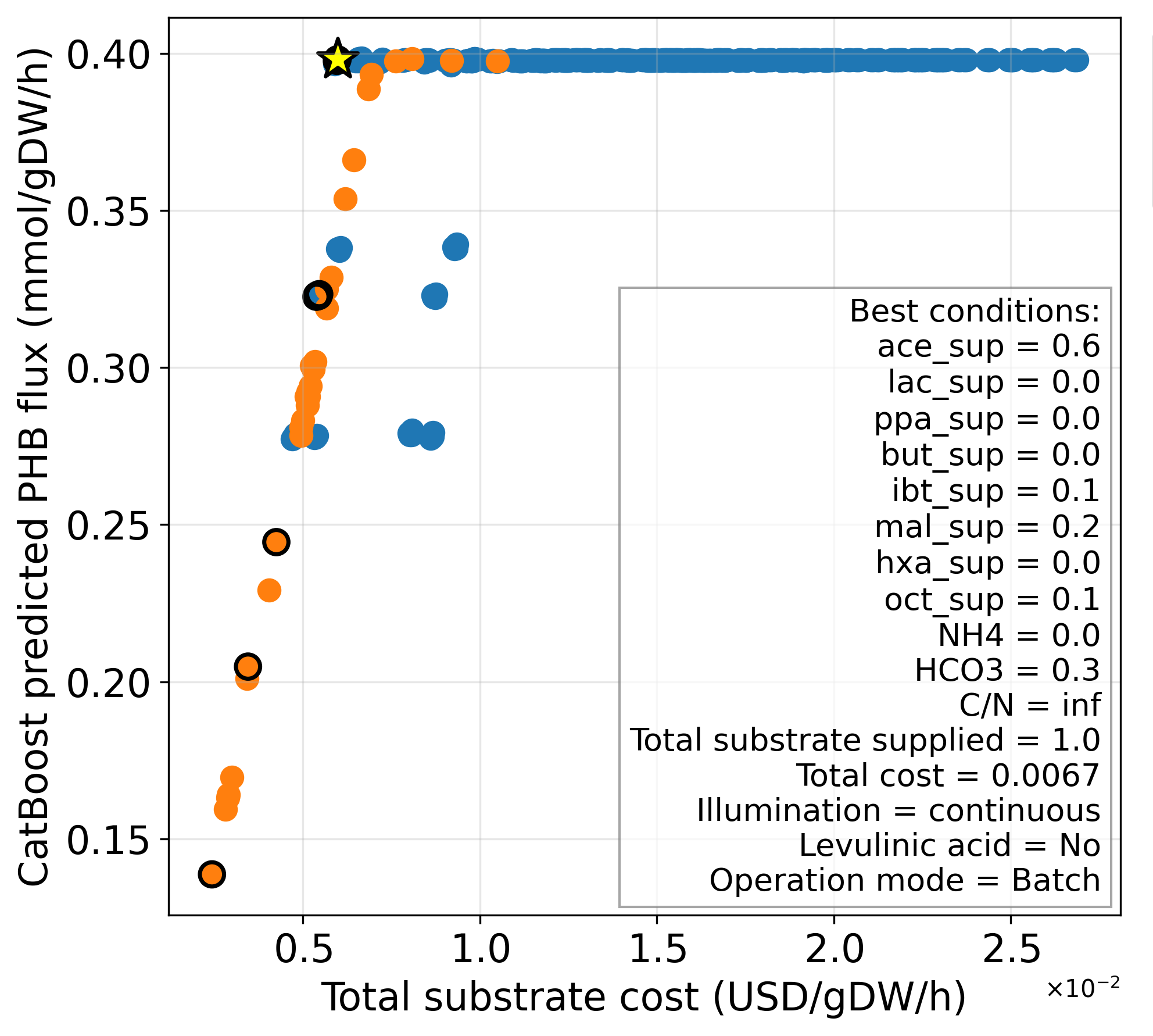

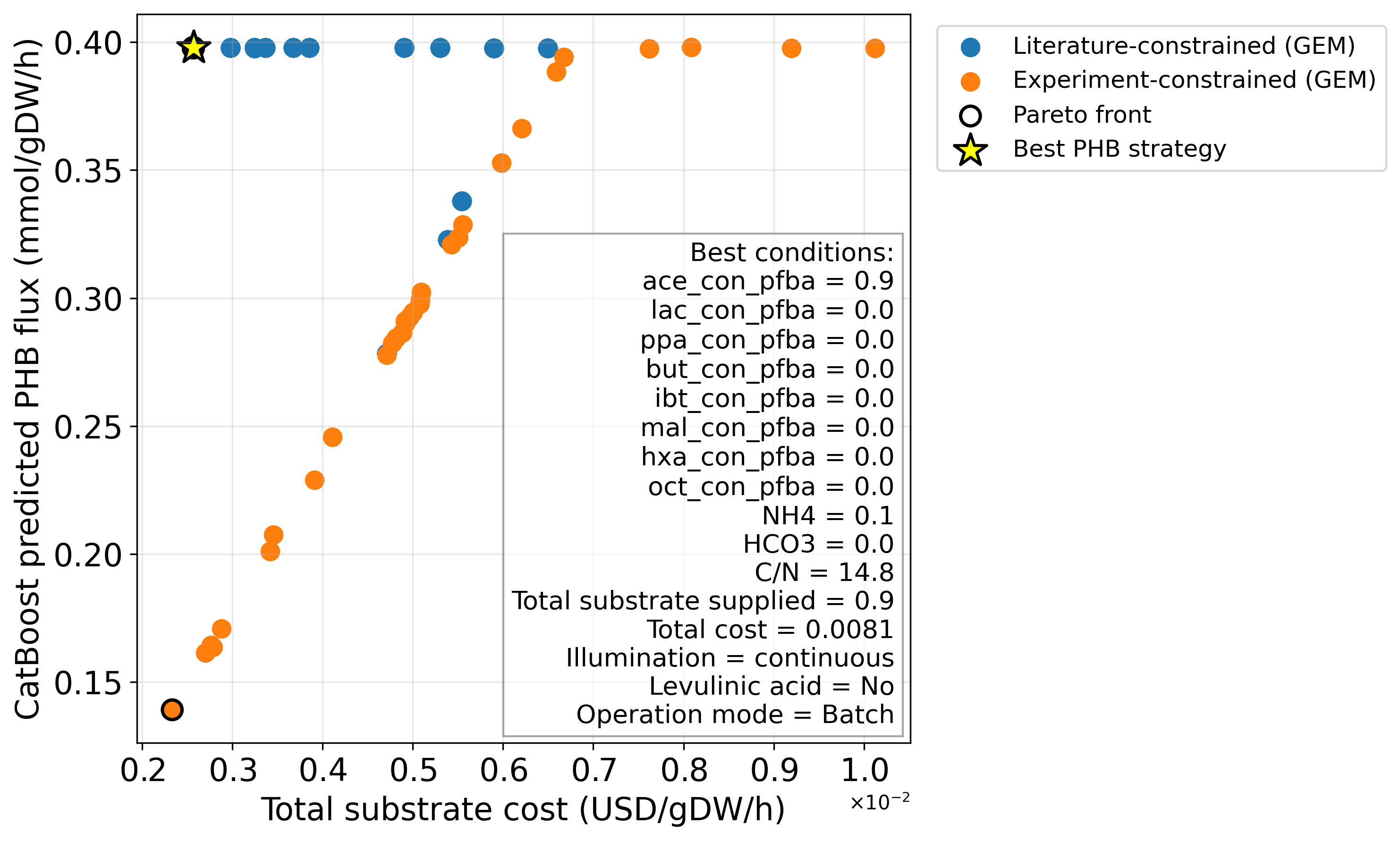

C)

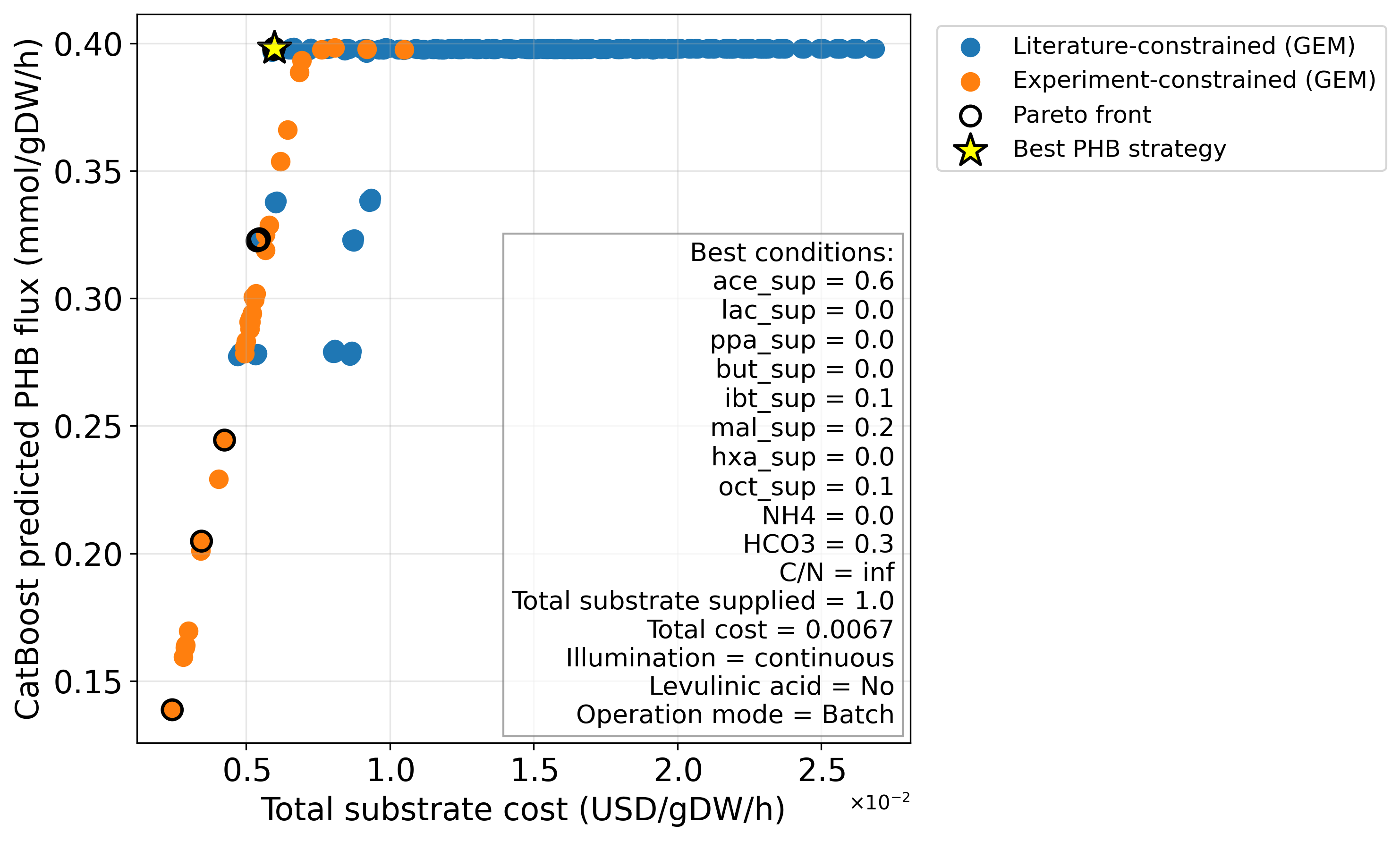

**Figure S1**. Results of CatBoost predicted and Pareto optimized PHB fluxes from parsimonious FBA (pFBA) with different datasets (PHB synthesis upper bond = 0.39 mmol/gDW-1/hr-1). A) Only supplied nutrient fluxes are presented to CatBoost, B) Only consumed nutrient fluxes are presented to CatBoost, C) Both supplied and consumed nutrient fluxes are presented to CatBoost. Yellow star represents the Pareto optimal solution with maximal PHB production.

Similarly to what it was done for standard FBA, conditions beyond the literature-based calculated metabolic limits of PHB synthesis flux (0.39 mmol gDW⁻¹ h⁻¹) were also explored. The PHB synthesis upper bound was left unconstrained during FBA. This scenario was evaluated *in silico* using the same workflow but increasing the PHB synthesis upper limit from 0.39 to 1000 mmol gDW⁻¹h⁻¹, the maximum allowable value in the GEM. The corresponding results are shown in Figure S2 (panels A–C). The three scenarios resulted in optimal TFA validation, meaning that they are thermodynamically feasible.

A)	B)

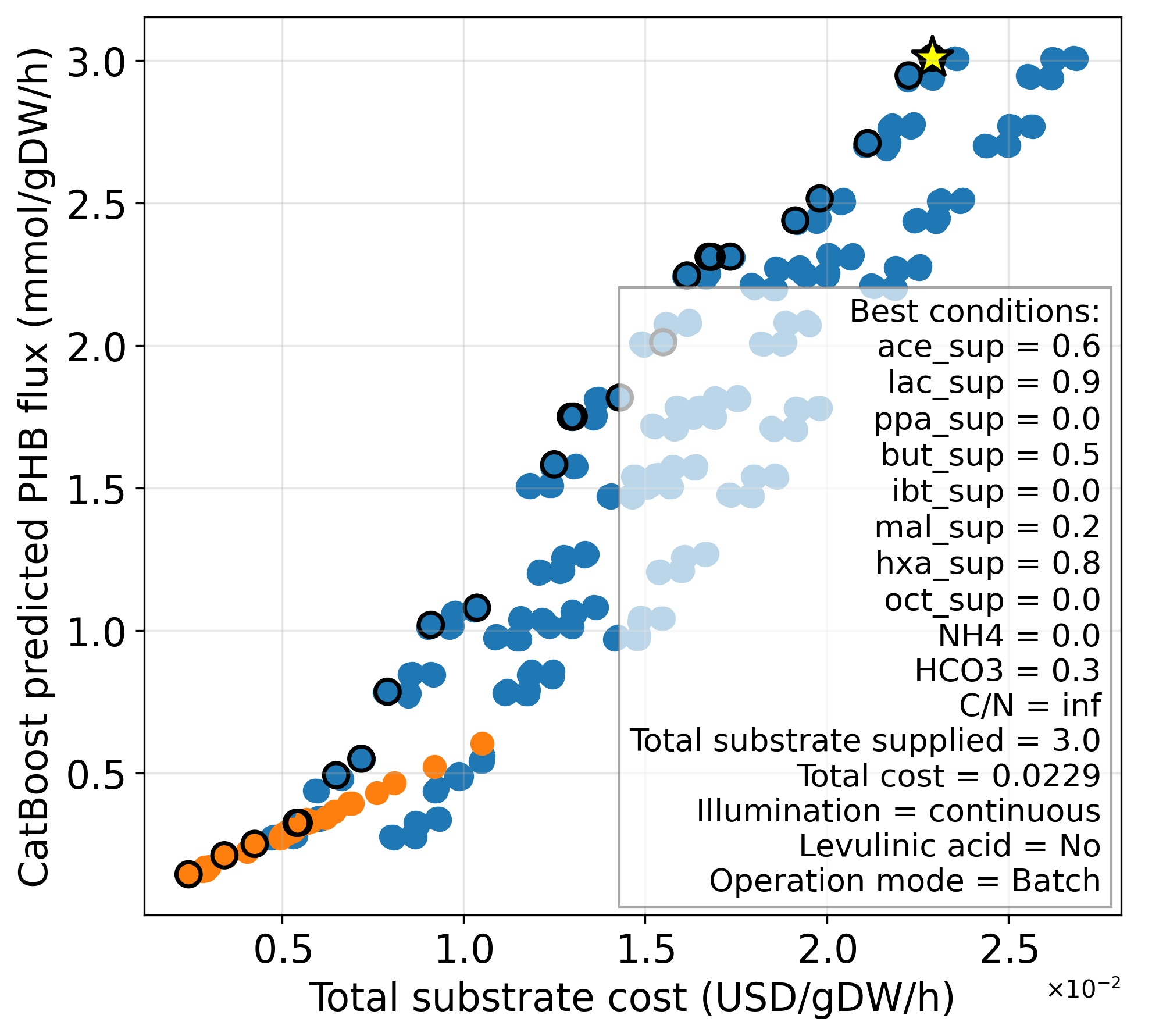

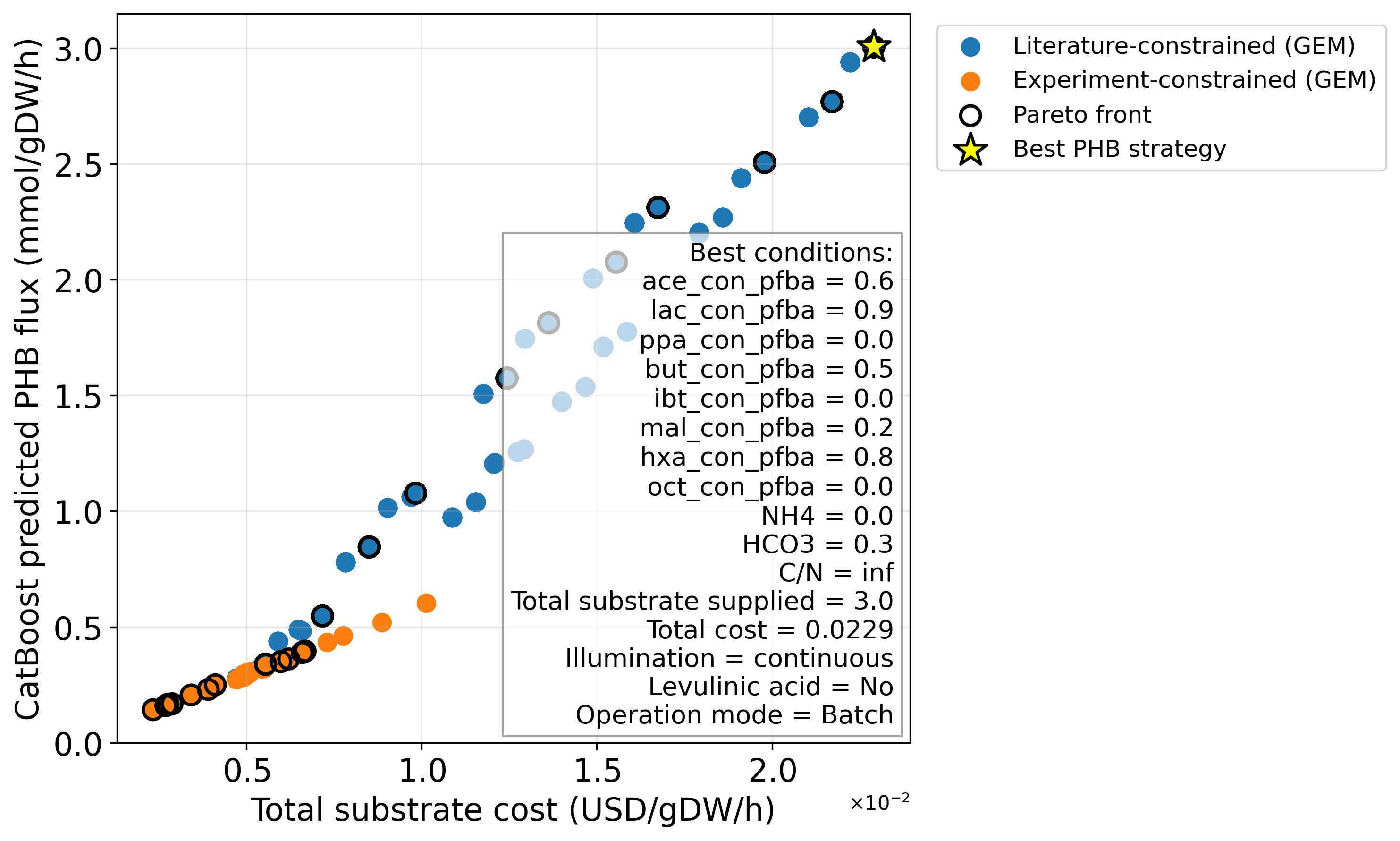

C)

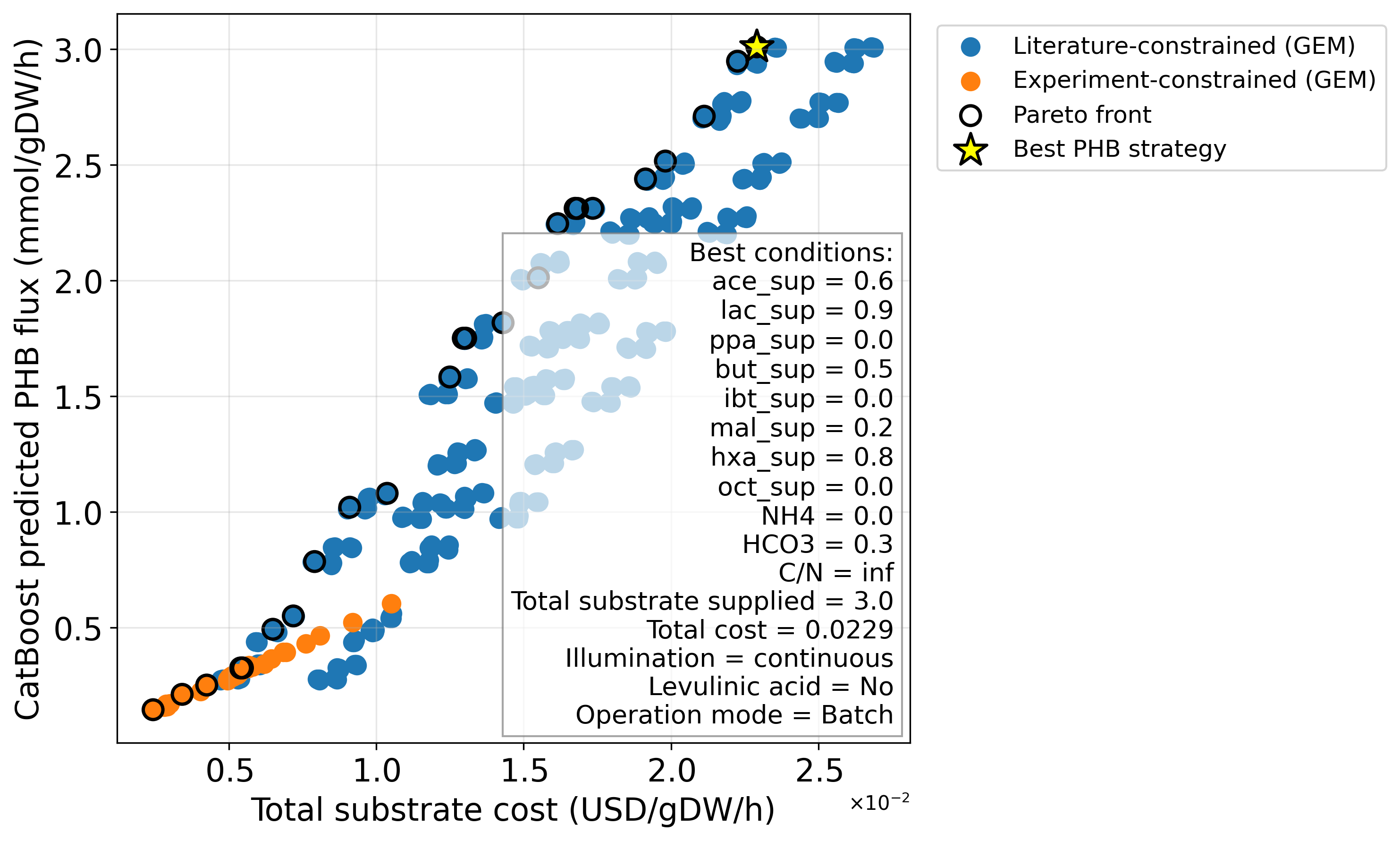

**Figure S2**. Results of CatBoost predicted and Pareto optimized PHB fluxes from parsimonious FBA (pFBA) with different datasets (PHB synthesis upper bond = 1000 mmol/gDW-1/hr-1). A) Only supplied nutrient fluxes are presented to CatBoost, B) Only consumed nutrient fluxes are presented to CatBoost, C) Both supplied and consumed nutrient fluxes are presented to CatBoost. Yellow star represents the Pareto optimal solution with maximal PHB production.

#### Top reactions and metabolic pathways using synthetic data for augmentation

Figure S3 shows the results of the flux ranking analysis for FBA and pFBA solutions, with an upper bound of 1000 mmol gDW^-1^h^-1^. Both profiles are dominated by strong light uptake (PHOA690um, EX_photon690_e) and active Photosystem I (PSIum, PSICSum), indicating that photosynthetic electron flow is central in both conditions. However, their downstream metabolic patterns diverge. For FBA, high flux through CBFCum, PGK, and GAPDi_nadp indicates a strongly active Calvin–Benson cycle, coupled to ATP synthesis (ATPSum, ATPM) and respiratory electron input (NDH_1_1_um_copy1). This pattern reflects a metabolism oriented toward carbon fixation and energy production, consistent with a growth- or biomass-driven state. In contrast, pFBA shows little Calvin cycle flux and instead exhibits increased activity in redox-balancing and central carbon redistribution reactions, including NADTRHD, FNOR_1, MDH, and acetyl-CoA–related steps (ACOAH, ACCOAC). This shift suggests reduced emphasis on carbon fixation and greater prioritization of intracellular redox control and precursor rerouting, consistent with a more reduced metabolic state that may favor storage or biosynthetic pathways rather than maximal growth.

A) FBA	B) pFBA

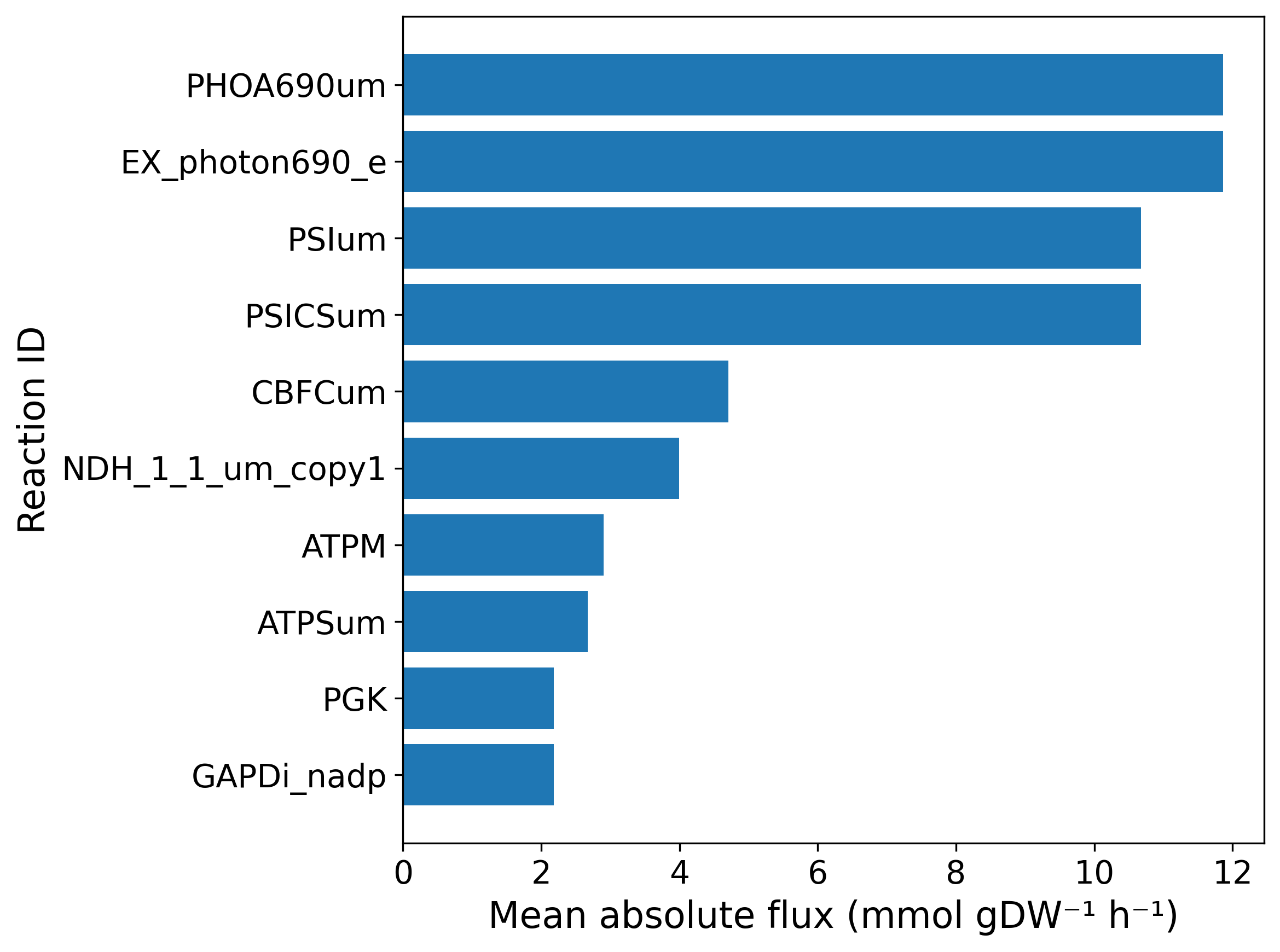

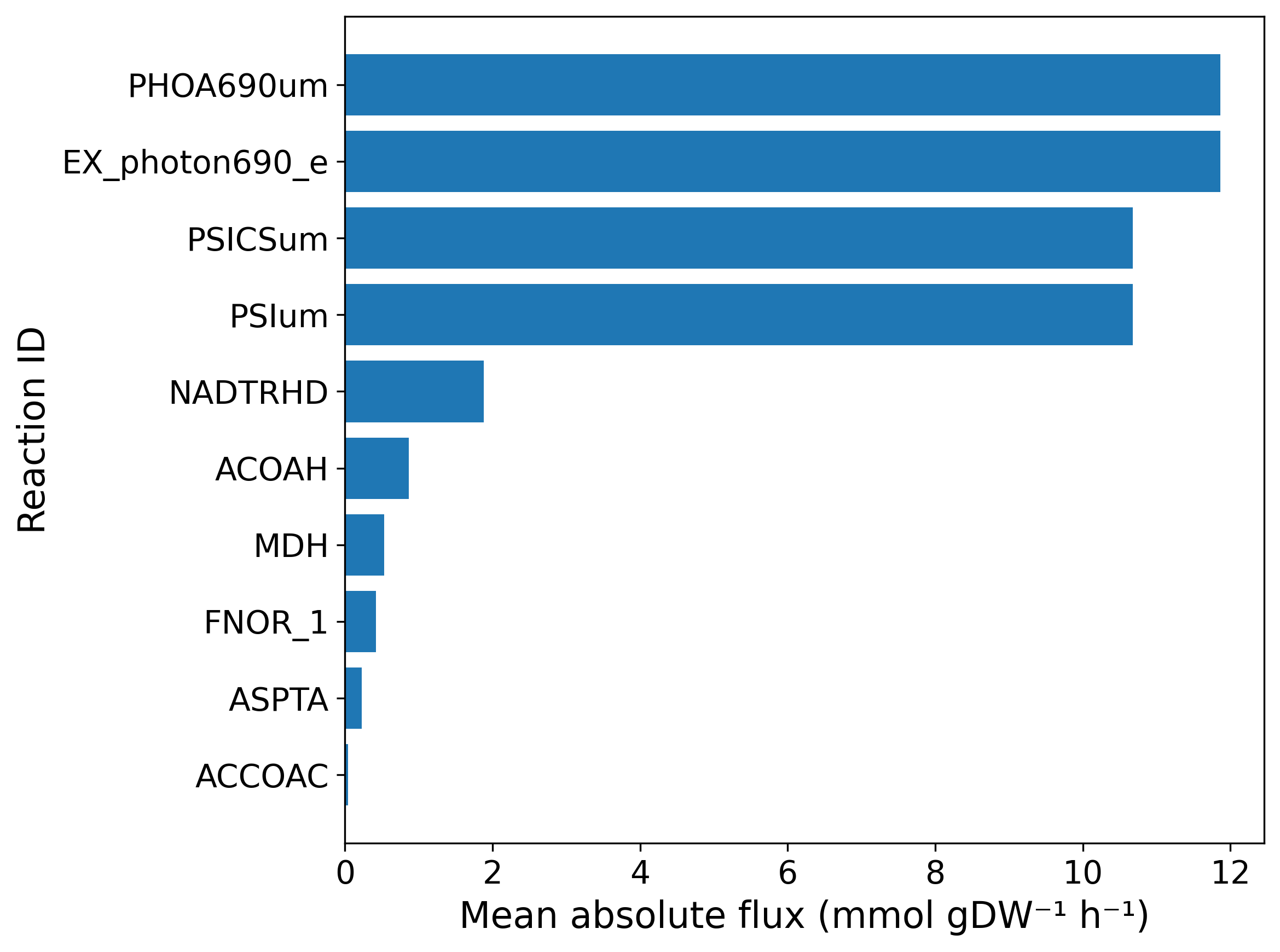

**Figure S3.** Top 10 reactions exhibiting the highest mean absolute fluxes across A) FBA and B) pFBA simulations with the PHB synthesis upper bound set to 1000 mmol gDW⁻¹ h⁻¹. IN both cases, the dominant reactions were primarily associated with photosynthetic processes, particularly photon absorption and light harvesting within the 680–700 nm wavelength range, nitrogen metabolism, amino acid metabolism, energy metabolism, and respiration pathways. In contrast, the pFBA solution exhibited a pronounced shift toward substrate-level metabolism and redox-efficient pathways.

#### SHAP plots for the unconstrained case

SHAP analysis was performed for the unconstrained FBA model (upper bound of 1000 mmol gDW⁻¹h⁻¹) to illustrate how feature contributions shift when the PHB synthesis reaction is not restricted by known metabolic limits (). Interestingly, the features with positive and negative effects on CatBoost model predictions are more strongly separated from each other in comparison to the case when the upper bound of1000 mmol gDW⁻¹h⁻¹ (see Figure 6). Low hexanoate, as well as intermediate to high values of cost and carbon supplied highly contributed to the model performance.

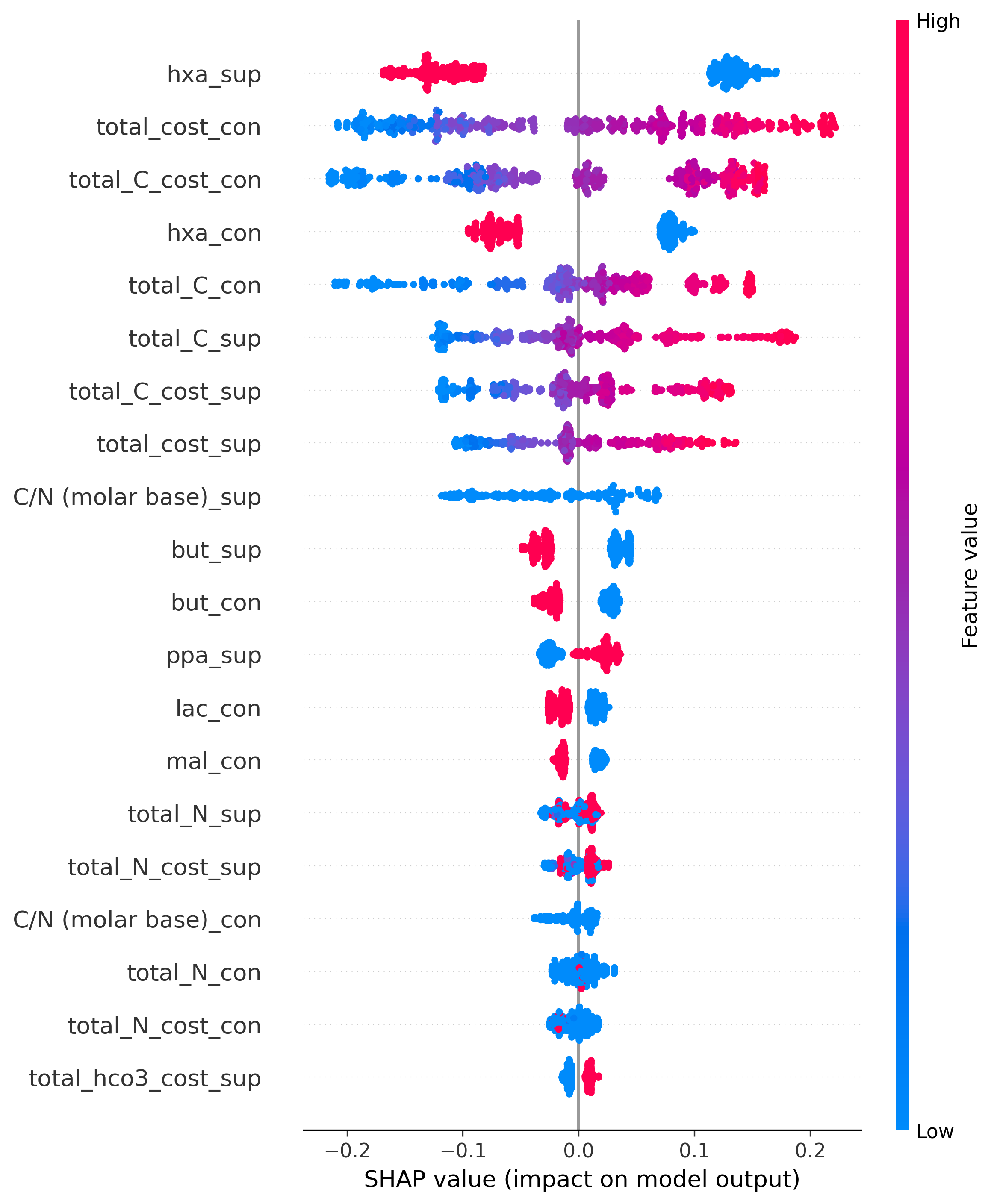

**Figure S4.** SHAP bee swarm plot of the CatBoost model constrained by the stoichiometric PHB synthesis limit (1000 mmol gDW⁻¹ h⁻¹) in FBA. By combining supplied (sup) and consumed (con) fluxes in FBA, the model captures the relationship between process conditions and metabolic capacity. Dot colors indicate feature magnitude, and the x‑axis shows their effect on the predicted PHB flux.

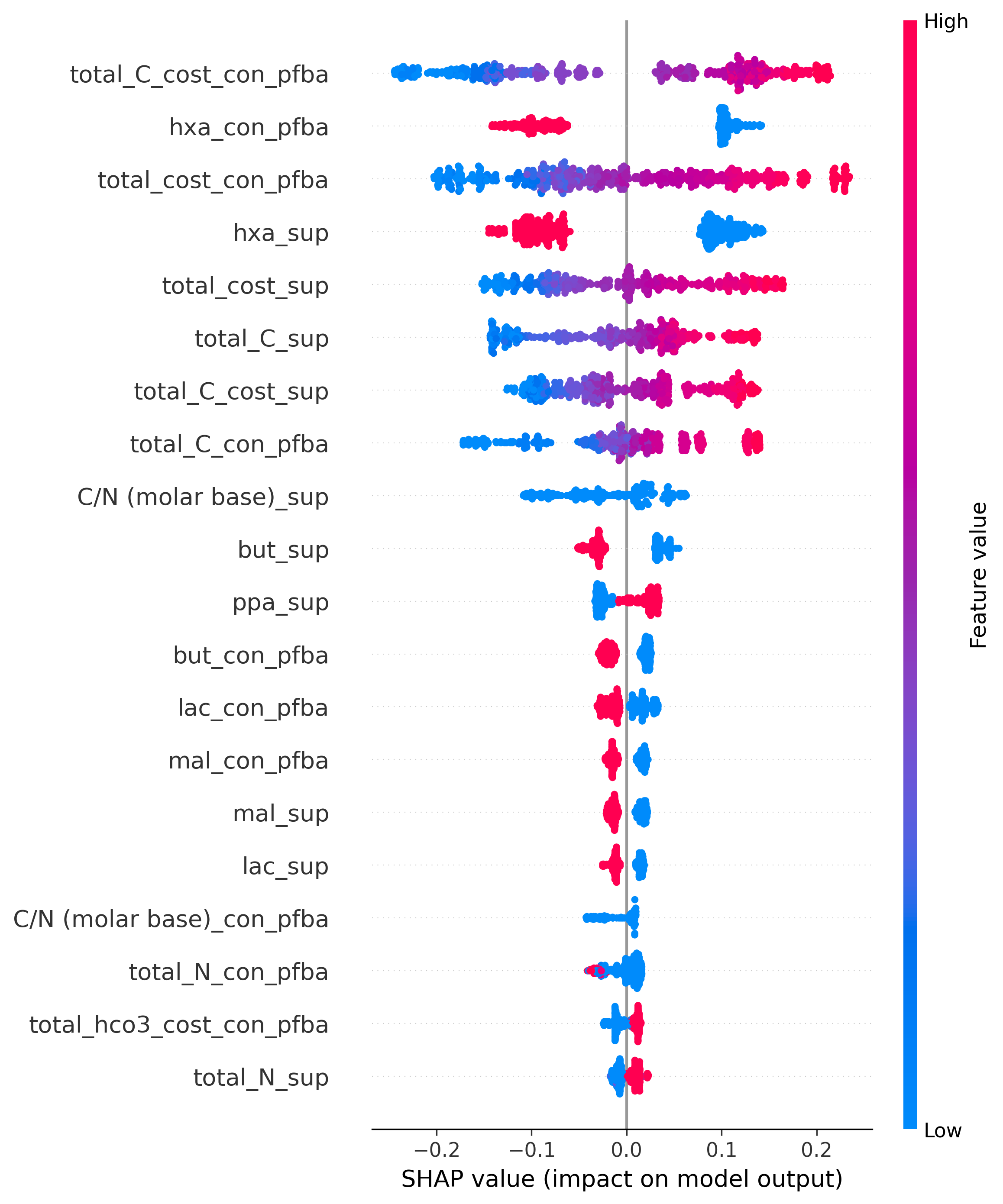

**Figure S5.** SHAP bee swarm plot of the CatBoost model constrained by the stoichiometric PHB synthesis limit (1000 mmol gDW⁻¹ h⁻¹). By combining supplied and consumed fluxes during pFBA, the model captures the relationship between process conditions and metabolic capacity. Dot colors indicate feature magnitude, and the x‑axis shows the effect on predicted PHB flux. Low hexanoate, along with intermediate to high values of cost and carbon supplied, contributed strongly to model performance.
